# Factors governing attachment of *Rhizobium leguminosarum* to legume roots

**DOI:** 10.1101/2022.11.21.517457

**Authors:** Jack D. Parsons, Clare R. Cocker, Alison K. East, Rachel M. Wheatley, Vinoy K. Ramachandran, Farnusch Kaschani, Markus Kaiser, Philip S. Poole

## Abstract

Primary attachment of rhizobia to host legume roots depends on pH and is the first physical interaction during nodulation. Genome-wide insertion sequencing, luminescence-based attachment assays and proteomic analysis demonstrate primary attachment of *Rhizobium leguminosarum* biovar *viciae* 3841 to *Pisum sativum* (pea) roots is more complex than previously thought. In total, 115 proteins are needed for initial attachment under one or more test conditions (acid, neutral or alkaline pH), with 22 required under all conditions. These include cell-surface filamentous hemagglutinin adhesin (RL4382) and its transporter (RL4381), transmembrane protein RL2400, RL3752 (PssA, glycosyl transferase) affecting capsular polysaccharide and transcriptional regulator RL4145 (PckR). RNASeq was used to determine targets of RL4145 (PckR) and regulator RL3453. The 54 proteins required for attachment at pH 7.0 were investigated for nodulation phenotypes. Glucomannan biosynthesis protein A (GmsA) is needed at pH 6.5 and pH 7.0. Membrane proteins DgkA and ImpA are required specifically at pH 6.5, and RpoZ at pH 7.5. Sonicated cell surface fractions inhibited root attachment at alkaline pH but no overlap between proteins identified by proteomic and INseq analysis, suggests there is no single rhicadhesin needed for alkaline attachment. Our results demonstrate the complexity of primary root attachment and diversity of mechanisms involved.

## Introduction

Rhizobia are diazotrophic soil Alphaproteobacteria which form nitrogen-fixing symbioses with host legume plants. Formation of these symbioses between rhizobia and their host legumes requires a complex and specific process of molecular cross-talk beginning in the rhizosphere (for a full review [1, 2]). In initial stages of symbiosis, host plants secrete a range of messenger molecules (including (iso)flavonoids) which induce rhizobial synthesis of Nodulation (Nod) factor (a decorated lipochitooligosaccharide), a specificity signal that has a role in triggering infection of the host plant and nodule organogenesis [1]. In the first physical interaction of the nascent symbiosis, rhizobia attach to the roots of the host legume. This attachment occurs in two distinct phases: primary attachment, defined as early-stage reversible interactions of single bacterial cells, and secondary attachment, defined as later-stage irreversible binding, often mediated by extracellular fibrils, and dependent on successful primary attachment [3]. Primary attachment of rhizobia to legume roots is necessary for subsequent successful formation of nitrogen-fixing symbioses, especially under the competitive conditions that exist in soil [4].

Much of the existing literature concerning primary root attachment in *Rhizobium*-legume symbioses describes a pH-dependent bacterial system based on a glucomannan/rhicadhesin duality (reviewed in [3, 5]). In this model, under mildly acidic soil conditions rhizobia use polarly-located glucomannan to bind to plant-root lectins [4, 6], while in mildly alkaline soil where lectins disassociate from roots, rhizobia make use of rhicadhesin (a putative 14kDa calcium-binding protein) whose encoding gene remains to be identified, to bind a (as yet uncharacterised) plant-root receptor for primary attachment [3, 5]. It is assumed that there is some overlap of these two systems under neutral soil conditions. However, the glucomannan/rhicadhesin hypothesis is likely to be incomplete as a model for primary attachment. Firstly, neither the rhicadhesin gene or its plant receptor have been identified. Secondly, while other factors have been shown to be important in primary attachment, they are often not taken into account. These include Van der Waals forces, electrostatic and hydrophobic interactions [3, 7, 8], as well as a range of molecular bacterial factors. Mutation of the extracellular polysaccharide (EPS) biosynthesis gene *pssA* in *Rhizobium leguminosarum* biovar *viciae* was demonstrated to disrupt ability to attach and form biofilms on inert surfaces, although it was not tested in any root attachment assays [4, 7]. Studies attempting to clone the rhicadhesin gene using an approach based on phage-display isolated several *Rhizobium*-adhering proteins (Raps) able to agglutinate cells, as well as promote biofilm formation and short-term attachment (four hour) to legume roots [9-11]. Furthermore, mutation of the regulator *praR* in *R. leguminosarum* bv. *viciae* was shown to enhance biofilm formation both on inert surfaces and legume roots (two hour assay), primarily through upregulation of Rap expression [12].

Therefore, although the glucomannan-rhicadhesin model is often used to explain primary root attachment in *Rhizobium*-legume symbioses, multiple other factors are likely to play a role. The rhizosphere is a complex and highly heterogeneous environment where both biotic and abiotic conditions vary [13-15]. With existing literature linking diverse proteins and processes to attachment [4, 7, 9-12], it suggests that the suite of factors necessary for primary attachment to plant roots may be more complex than previously thought.

Existing tools for the investigation of attachment, while providing useful approaches for characterising novel factors, are proving to be rate-limiting in fully characterising primary attachment, in part due to the wide range of different approaches. Primary attachment assays, ranging from one to four hours in duration, have used wild type (WT) or mutant rhizobia and phase contrast microscopy to count bacteria attached to roots, or vortexing/sonication of roots to remove bacteria before plating them on growth media for colony enumeration [11, 16-18]. However, these methods are low-throughput, may suffer from ‘region-bias’ when counting attached bacteria using microscopy and will result in the loss of spatial attachment data if bacteria are removed before counting. Frederix *et al*. [12] developed a scintillation-based attachment assay using radiolabelled rhizobia and, subsequently, a luminescence (Lux)-based attachment assay by making use of rhizobia carrying a stably inherited plasmid encoding the *luxCDABE* operon. Each of the above methods made use of excised root sections, which introduces an unnatural source of root exudation from the wound site, potentially affecting attachment. Furthermore, none were tested for their suitability in conducting primary attachment assays across a range of pHs. Initial reports of the effects of pH on root attachment come from a variety of different methods, some growing the bacteria at one pH and assessing attachment at another [4, 6]

In this study, we have developed a standard one hour primary attachment assay and used it to determine genetic factors important for primary root attachment of *Rhizobium leguminosarum* biovar *viciae* 3841 (Rlv3841) [19] to roots of the host legume *Pisum sativum* (pea). Using *mariner* transposon-based Insertion Sequencing (INSeq) [20], attachment experiments were performed under acid, neutral and alkaline plant growth conditions. INSeq enables a holistic study of gene fitness under conditions of interest by using high-throughput sequencing of a bacterial insertion mutant library isolated from different environmental conditions (in this case from pea roots at acid, neutral and alkaline pH), combined with hidden Markov model (HMM) analysis [21-23]. INSeq results were compared with data from whole-root Lux-based bacterial attachment assays performed under the same three pH conditions. We identified key genes for primary attachment under all conditions tested (pH 6.5, 7.0 and 7.5), as well as at specific pHs. By combining INSeq, Lux-based attachment data and a proteomics approach, we attempted to identify the rhicadhesin gene. A model for the primary attachment of Rlv3841 to pea roots is presented which illustrates that key regulator genes are required to facilitate the interactions involved.

## Materials and methods

### Strains, plasmids and culture conditions

All strains and plasmids used are listed in Supplementary Table S1. Rhizobial strains were grown on universal minimal salts (UMS) medium at 28°C as previously described [22], supplemented with 30 mM sodium pyruvate, 10 mM ammonium chloride and adjusted to pH 6.5, pH 7.0 or pH 7.5. For solid media 1% w/v agar was added. All *Escherichia coli* strains were grown at 37 °C in Lennox (L)-broth or on L-agar [24]. Antibiotics were used at the following concentrations (μg ml^-1^), unless otherwise stated: neomycin, 80; streptomycin, 500; tetracycline, 5. The donor *E. coli* strain SM10*λpir* (carrying the *mariner* transposon INSeq pSAM_Rl vector [20]) was supplemented with 50 μg ml^-1^ neomycin to select for the plasmid and 100 μg ml^-1^ ampicillin to select for *E. coli*. Following recovery of attached bacteria in the INSeq attachment assays, bacteria were regrown for 12 h in tryptone-yeast extract (TY) [24], prior to DNA extraction.

*Rhizobium* mutant strains were made using single-crossover integration pK19mob-mutagenesis by first making a clone of the gene using PCR [25]. Integration of pK19 into the correct gene was confirmed by PCR-mapping (primers listed in Supplementary Table S2). For all Lux-based attachment assays [12], *Rhizobium* strains were labelled by introduction of pIJ11282 (plasmid with *luxCDABE* constitutively expressed from *nptII* promoter and maintained in rhizobia [12]). Plasmid pIJ11282 was introduced into rhizobial strains according to the method of Figurski and Helinski [26].

### *Mariner* transposon library construction

The *mariner* transposon library was constructed as previously described by Wheatley et al. [22]. Briefly, donor *E. coli* were grown in L-broth overnight and Rlv3841 was grown on a TY agar slope. Cultures were pelleted, resuspended in TY and pooled. Cells were further pelleted and resuspended in TY before spotting cell suspensions on nitrocellulose filters on TY agar plates and incubation at 28°C overnight. Filters were resuspended in UMS with 15% glycerol before enumeration of transposon insertions in Rlv3841 on TY agar supplemented with streptomycin and neomycin and pooling of mutants.

### Buffering capacity of vermiculite

Rooting solution (25 ml) (consisting of 1 mM CaCl_2_.2H_2_O, 100 μM KCl, 800 μM MgSO_4_.7H_2_O, 10 μM Fe EDTA, 35 μM H_3_BO_3_, 9 μM MnCl_2_.4H_2_O, 0.8 μM ZnCl_2_, 0.5 μM Na_2_MoO_4_.2H_2_O, 0.3 μM CuSO_4_.5H_2_O, 4 mM KH_2_PO_4_, 4 mM Na_2_HPO_4_), was adjusted to pH 6.5, 7.0 or 7.5 using hydrochloric acid or sodium hydroxide solution and added to 2.5 g fine vermiculite (Sinclair) in 50-ml Falcon tubes, in triplicate. Tubes were rotated gently at 20 rpm and the pH of rooting solution measured using a pH meter (Hanna) at time intervals up to 72 h. Although slight pH increases were seen over time, the pH never increased by more than 0.16 above the initial pH (Supplementary Fig. S1).

### Plant growth and root attachment assays

Seeds of *Pisum sativum* (var. Avola) were sterilised in a 5% sodium hypochlorite solution for 5 min before washing six times with distilled water. Washed fine vermiculite (30 g) was placed in 100ml-boiling tubes, together with 25 ml of rooting solution (adjusted to pH 6.5, pH 7.0 or pH 7.5, depending on the experiment). Boiling tubes were sterilised, and pea seeds planted under sterile conditions were grown for 7 d. Three days prior to the experiment, Rlv3841-based strains (*mariner* transposon library or individual Lux-labelled strains) were streaked on UMS agar slopes. For INSeq experiments, all slopes were at pH 7.0. For Lux-based attachment assays, the pH of the slopes was adjusted to the test pH using hydrochloric acid or sodium hydroxide solution. For both INSeq experiments and Lux-based assays the same procedure to assess attachment was followed: bacteria were resuspended in 15 mM MES/HEPES (pH adjusted as appropriate) to an OD_600_ of 0.1. Plants were removed from vermiculite, washed in 15 mM MES/HEPES (of the correct pH), and their roots submerged in 50 ml of the bacterial suspension for 1 h, with gentle (20 rpm) agitation. After 1 h, roots were removed and washed by dipping in 15 mM MES/HEPES (of the correct pH). For INSeq experiments, roots were excised from the seed and vortexed (Heidolph Multi Reax) at maximum speed for 10 min in 50 ml-Falcon tubes with 20 ml 15 mM MES/HEPES. Roots were discarded, and the pooled liquid filtered through 3 layers of sterile muslin cloth before re-growing bacteria in 50 ml TY media at 28°C for 12 h (with shaking at 180 rpm). This step was included to boost bacterial DNA, helping to minimise contamination by plant material. DNA was then isolated using a Qiagen DNeasy Blood and Tissue Kit for Gram-negative bacteria according to the manufacturer’s protocol with the modifications described in [22]. For the input library, DNA was isolated directly from 50 ml of OD_600_ = 0.1 resuspension (i.e., the bacterial suspension into roots were dipped) as described above. INSeq was performed in triplicate for each test condition (root attachment at acid, neutral and alkaline) and for the input library.

For Lux-based bacterial attachment assays [12], weighed pea roots (*n* = 10) were imaged using a NightOWL LB 983 imaging system (Berthold). Root-attached luminescence was normalised using inoculum luminescence, measured in triplicate using a GloMax plate reader (Promega). Bacterial attachment to roots is expressed as Relative Light Units (RLU)/g of root [12]. To assess the effect of a crude preparation of adhesin, sterilised peas were germinated for 5 d on water agar. Roots were excised from seed and seedlings, and preincubated with the crude adhesin fraction (600 μg protein) for 1 h before washing by dipping in 15 mM MES/HEPES before bacterial attachment assessed in Lux-based assays.

### Library preparation and sequencing

Following DNA isolation, transposon tags were prepared for DNA sequencing as described in [22]. Briefly, linear PCR products were amplified using BioSAM primers [20] with 1000 ng DNA (see Supplementary Table S2 for primer and adaptor sequences). Biotinylated linear PCR products were bound to Pierce streptavidin magnetic beads (Thermo Scientific) and enzymatic library preparation performed. A custom INSeq library adaptor was used for adaptor ligation and final PCR-amplification performed using one of 12 different barcoding primers (Supplementary Table S2). The final sequencing template for each group (187-bp in length) was purified using SizeSelect II agarose gels (Invitrogen) and analysed using a Bioanalyzer High Sensitivity DNA chip (Agilent Technologies). Libraries were diluted to 100 pM and pooled. Libraries were subject to automatic template preparation using an Ion Chef (Life Technologies) and DNA sequencing was performed on an Ion Proton Sequencer (ThermoFisher Scientific).

### Transposon insertion analysis and HMM

Sequencing reads were analysed on a Linux server using a custom-written Perl script as previously described [20, 22] with ‘cutadapt’ [27], the Bowtie short read aligner [28] and the Tn-HMM Python module [29]. HMM analysis was used to assign each gene to one of four categories: essential (ES), growth-deficient (GD), neutral (NE), and growth-advantage (GA) [29], and used for classification of genes in the input library. However, for attachment experiments the following were adopted: essential (ES), defective (DE), neutral (NE) or advantaged (AD), as used by Wheatley at al. [23]. HMM categories were averaged across the three biological replicates per test condition. Manual curation of those genes lacking a consensus classification was performed. This was achieved by visualising INSEQ reads mapped to the Rlv3841 genome using the Integrative Genomics Viewer platform and assessing TA insertion profiles between input and experimental samples. Visualisation enabled direct comparison of numbers of TA site insertions mapping in and around gene loci between input and experimental samples. This provided a further control for assessing fitness impacts under input and experimental conditions, enabling an indicative consensus state call to be assigned. This applied to 27 genes, listed in Datasheet S1 (marked mc (manually curated)). Following removal of genes which have no TA sites (159), non-NE in the input library (compromised before root attachment investigated) (947), non-NE for growth in TY [20] (compromised after isolation from roots) (156) and pseudogenes (9), a total of 5,986 genes were assessed for their role in primary attachment (Datasheet S1).

### Crude adhesin isolation and proteomics

Rlv3841 was grown overnight at 28°C in 250 ml UMS with shaking at 180 rpm. Cell fractionation and isolation of the crude adhesin fraction was carried out as previously described [30]. Briefly, cells were harvested by centrifugation at OD_600_= 0.7, and the cell pellet washed and resuspended in 25mM phosphate buffer. Cells were sheared using a 705 Sonic Dismembrator (Fisher Scientific) at an amplitude of 5. The suspension was centrifuged at 12,000 x g for 10 min at 4°C, and the resulting supernatant then centrifuged at 100,000 x g (Beckmann TL-100 ultracentrifuge) for 2 hr at 4°C. The supernatant obtained at this stage formed the crude adhesin fraction. Prior to root attachment assays, the concentration of protein in the crude adhesin fraction was determined using a Pierce Bicinchoninic Acid (BCA) assay (Thermo Scientific) according to manufacturer’s instructions. Root attachment assays were performed following pre-incubation for 1 h with 600 μg crude adhesin protein. Separation of crude adhesin was achieved using Nu-PAGE Bis-Tris gels (Thermo Scientific) according to manufacturer’s instructions. For protein visualisation, SYPRO Ruby staining (Sigma Aldrich) was used according to the manufacturer’s ‘rapid’ protocol. Imaging was carried out using a BioRad ChemiDoc XRS+ imaging system with ImageLab (BioRad). For proteomic analysis, bands of interest were excised and subjected to liquid chromatography-mass spectrometry (LC-MS). Calcium-binding potential of candidate peptides was assessed through the IonCom server tool [31].

### In-gel digestion (IGD) with trypsin

Gel bands of interest were cut out of the SDS Page gel and subjected to IGD following a published protocol [32]. The acidified tryptic digests were finally desalted on home-made 2 disc C18 StageTips as described [33]. After elution from the StageTips, samples were dried using a vacuum concentrator (Eppendorf) and the peptides were taken up in 10 μL 0.1 % formic acid solution.

### LC-MS/MS settings

Experiments were performed on an Orbitrap Elite instrument (Thermo) [34] that was coupled to an EASY-nLC 1000 liquid chromatography (LC) system (Thermo). The LC was operated in the one-column mode. The analytical column was a fused silica capillary (75 μm × 35 cm) with an integrated PicoFrit emitter (New Objective) packed in-house with Reprosil-Pur 120 C18-AQ 1.9 μm resin (Dr. Maisch). The analytical column was encased by a column oven (PRSO-V1; Sonation) and attached to a nanospray flex ion source (Thermo). The column oven temperature was adjusted to 45 °C during data acquisition. The LC was equipped with two mobile phases: solvent A (0.1% formic acid, FA, in water) and solvent B (0.1% FA in acetonitrile, ACN). All solvents were of UPLC grade (Sigma-Aldrich). Peptides were directly loaded onto the analytical column with a maximum flow rate that would not exceed the set pressure limit of 980 bar (usually around 0.6 – 1.0 μL/min). Peptides from in-gel digests (IGD) were subsequently separated on the analytical column by running a 70 min gradient of solvent A and solvent B (start with 7% B; gradient 7% to 35% B for 60 min; gradient 35% to 80% B for 5 min and 80% B for 5 min) at a flow rate of 300 nl/min. The mass spectrometer was operated using Xcalibur software (version 2.2 SP1.48). The mass spectrometer was set in the positive ion mode. Precursor ion scanning was performed in the Orbitrap analyzer (FTMS; Fourier Transform Mass Spectrometry) in the scan range of m/z 300-1500 (IGD) or 1800 (ISD) and at a resolution of 60000 with the internal lock mass option turned on (lock mass was 445.120025 m/z, polysiloxane) [35]. Product ion spectra were recorded in a data dependent fashion in the ion trap (ITMS) in a variable scan range and at a rapid scan rate (Wideband activation was turned on). The ionization potential (spray voltage) was set to 1.8 kV. Peptides were analyzed using a repeating cycle consisting of a full precursor ion scan (3.0 × 10^6^ ions or 50 ms) followed by 12 product ion scans (1.0 × 10^4^ ions or 80 ms) where peptides are isolated based on their intensity in the full survey scan (threshold of 500 counts) for tandem mass spectrum (MS2) generation that permits peptide sequencing and identification. Collision induced dissociation (CID) energy was set to 35% for the generation of MS2 spectra. During MS2 data acquisition dynamic ion exclusion was set to 120 seconds with a maximum list of excluded ions consisting of 500 members and a repeat count of one. Ion injection time prediction, preview mode for the FTMS, monoisotopic precursor selection and charge state screening were enabled. Only charge states higher than 1 were considered for fragmentation.

### Peptide and Protein Identification using MaxQuant

RAW spectra were submitted to an Andromeda [36] search in MaxQuant (1.5.3.30 or) using the default settings [37]. Label-free quantification and match-between-runs was activated [38]. The Uniprot reference Database for *R. leguminosarum* bv. *viciae* (strain 3841) was used for the search (UP000006575_216596.fasta; 7093 entries; downloaded 06.12.2016). All searches included a contaminants database search (as implemented in MaxQuant, 245 entries). The contaminants database contains known MS contaminants and was included to estimate the level of contamination. Andromeda searches allowed oxidation of methionine residues (16 Da) and acetylation of the protein N-terminus (42 Da) as dynamic modifications and the static modification of cysteine (57 Da, alkylation with iodoacetamide). Enzyme specificity was set to “Trypsin/P” with two missed cleavages allowed. The instrument type in Andromeda searches was set to Orbitrap and the precursor mass tolerance was set to ±20 ppm (first search) and ±4.5 ppm (main search). The MS/MS match tolerance was set to ±0.5 Da. The peptide spectrum match FDR and the protein FDR were set to 0.01 (based on target-decoy approach). Minimum peptide length was 7 amino acids. For protein quantification unique and razor peptides were allowed. Modified peptides were allowed for quantification. The minimum score for modified peptides was 40. Label-free protein quantification was switched on, and unique and razor peptides were considered for quantification with a minimum ratio count of 2. Retention times were recalibrated based on the built-in nonlinear time-rescaling algorithm. MS/MS identifications were transferred between LC-MS/MS runs with the “match between runs” option in which the maximal match time window was set to 0.7 min and the alignment time window set to 20 min. The quantification is based on the “value at maximum” of the extracted ion current. At least two quantitation events were required for a quantifiable protein. Further analysis and filtering of the results was done in Perseus [39]. Comparison of protein group quantities (relative quantification) between different MS runs is based solely on the LFQ’s as calculated by the MaxQuant MaxLFQ algorithm [38].

### Data availability

The mass spectrometry proteomics data for the on-bead digestions have been deposited to the ProteomeXchange Consortium via the PRIDE [40] partner repository (https://www.ebi.ac.uk/pride/archive/) with the dataset identifier PXD029089. During the review process, the data can be accessed via a reviewer account (**Username:**; **Password:** yBcwsy6Z).

### RNASeq

Strains OPS1907, OPS1908 and RU4062 (pK19 mutants in RL3453, RL4145 (*pckR*) and pRL100162 (*nifH*), respectively) (Table S1) were grown for three days at 28°C on UMS agar slopes. Bacteria were resuspended in 1:1 UMS and RNAlater. RNA extraction was carried out using a RNeasy mini kit (Qiagen) and genomic DNA depleted using a TURBO DNA-free kit (Ambion). Sequencing was carried out by Novogene using the Illumina NovoSeq 6000. Analysis was performed as described previously [41]. Data has been uploaded to the NCBI SRA database with the accession number PRJNA811156.

### General computing

Geneious R10 was used for primer design and local sequence alignment [42]. Global nucleotide and protein sequence alignments were carried out using BLASTn and BLASTp (NCBI) [43]. Protein-protein interaction networks were predicted and visualized using STRING [44]. Cellular protein localization was predicted using pSORTb v 3.0.2 [45]. In bacterial attachment assays, luminescence was evaluated using IndiGO software (Berthold) and subject to statistical testing in GraphPad Prism 8. Data handling was largely in MS Excel and all graphs were generated using GraphPad Prism 8. RpoH1 sequence searches were carried out using FIMO from MEME suite of software [46].

## Results

### Lux-based assay for primary attachment of *R. leguminosarum* to pea roots

A Lux-based assay was developed where primary attachment is defined as the attachment of bacteria to pea roots after one hour of incubation. We define this as primary attachment because we saw no statistically significant difference between the number of bacteria recovered from pea roots by reversible attachment (vortexing roots) and irreversible attachment (vortexing and grinding of roots) (Supplementary Fig. S2).

Using the Lux-based attachment assay (bacteria constitutively labelled with *luxABCX*) [12], primary attachment of wild-type (WT) Rlv3841 to pea roots at pH 7.0 was approx. 1.5 × 10^5^ RLU /g of root (Fig. 1). Results for previously reported strains with mutations in genes shown to affect primary attachment are compared with WT in Fig. 1. A *pssA* mutant shows approx. 5 × 10^4^ RLU/g of root, which is significantly different from WT (*p* < 0.01) (Fig. 1). The *pssA* (RL3752) gene encodes a glycosyl transferase involved in acidic EPS biosynthesis and a mutant in *pssA* lacks both EPS and capsular polysaccharide [4]. In agreement with previous results a strain mutated in two genes each encoding a *Rhizobium* adherence protein (Rap) (*rapA2/rapC* double mutant) [12] shows an even lower level of attachment to pea roots, approx. 3 × 10^4^ RLU/g of root (*p* < 0.005) (Fig. 1).

**Fig. 1.**
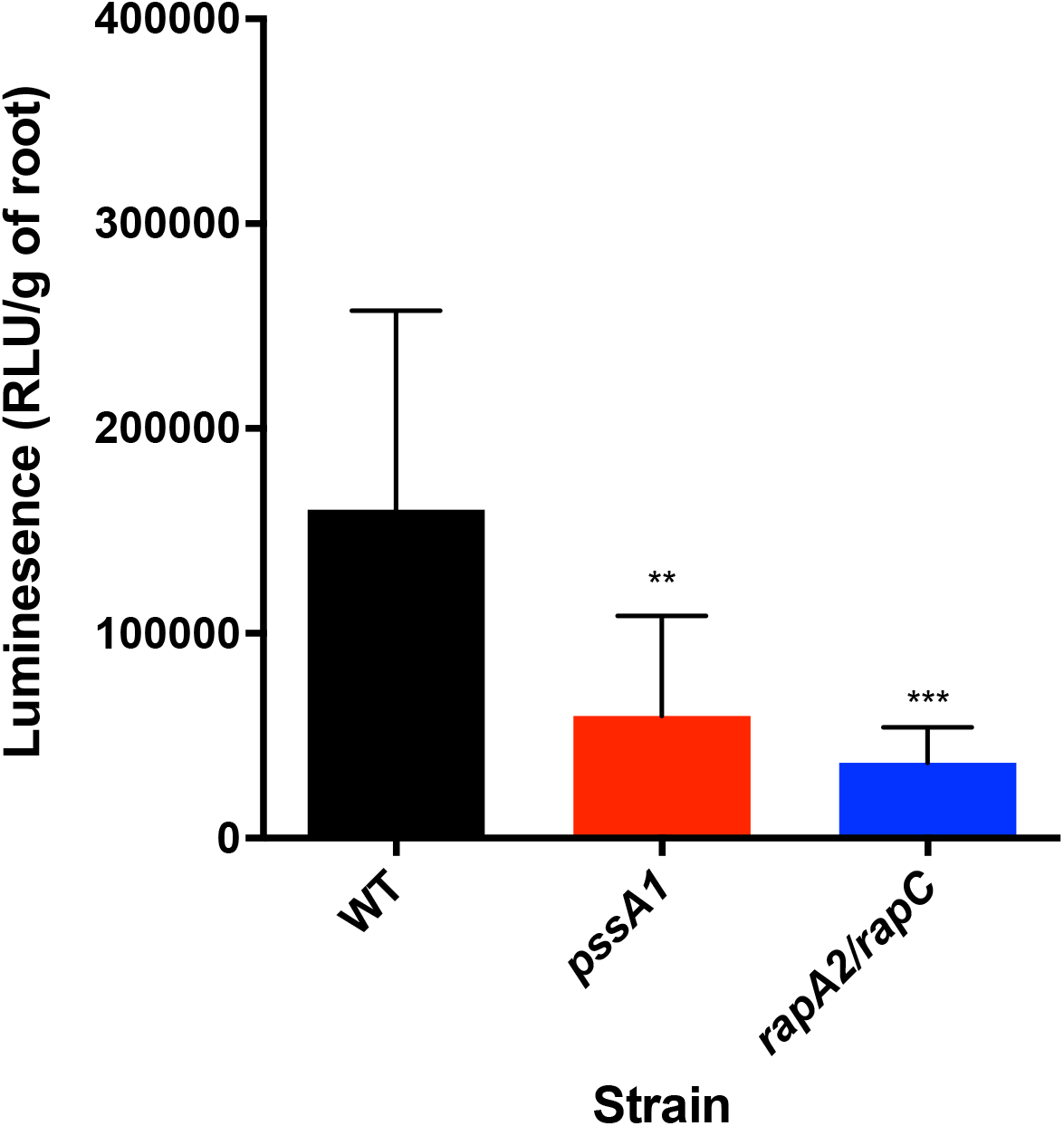
Primary attachment to pea roots of Rlv3841 (WT) and strains mutated in *pssA1* (RL3752) and *rapA2/rapC* (pRL100451/RL3074) at pH 7.0. Measured using the Lux-based whole root attachment assay where each strain is labelled by introduction of a plasmid constitutively expressing Lux. WT is shown in black, *pssA1* in red and *rapA2/rapC* in blue. Luminescence (RLU/g of root) shows bacterial attachment after 1 h. *n* = 10. Data are expressed as the mean ± SD. **= *p* < 0.01 and ***= *p* < 0.005 in comparison to WT using Student’s t-test.

Primary attachment of WT was not statistically different at any pH from 6.5-7.5 (Fig. 2). A *gmsA* strain, lacking glucomannan [4], had reduced attachment compared to WT (*p* < 0.001) at pH 6.5 and pH 7.0, but not at pH 7.5 (Fig. 2). Therefore, 1 h Lux-based assays for primary attachment (Figs. 1 & 2) agree with previous results for key mutants affected in primary adhesion.

**Fig. 2.**
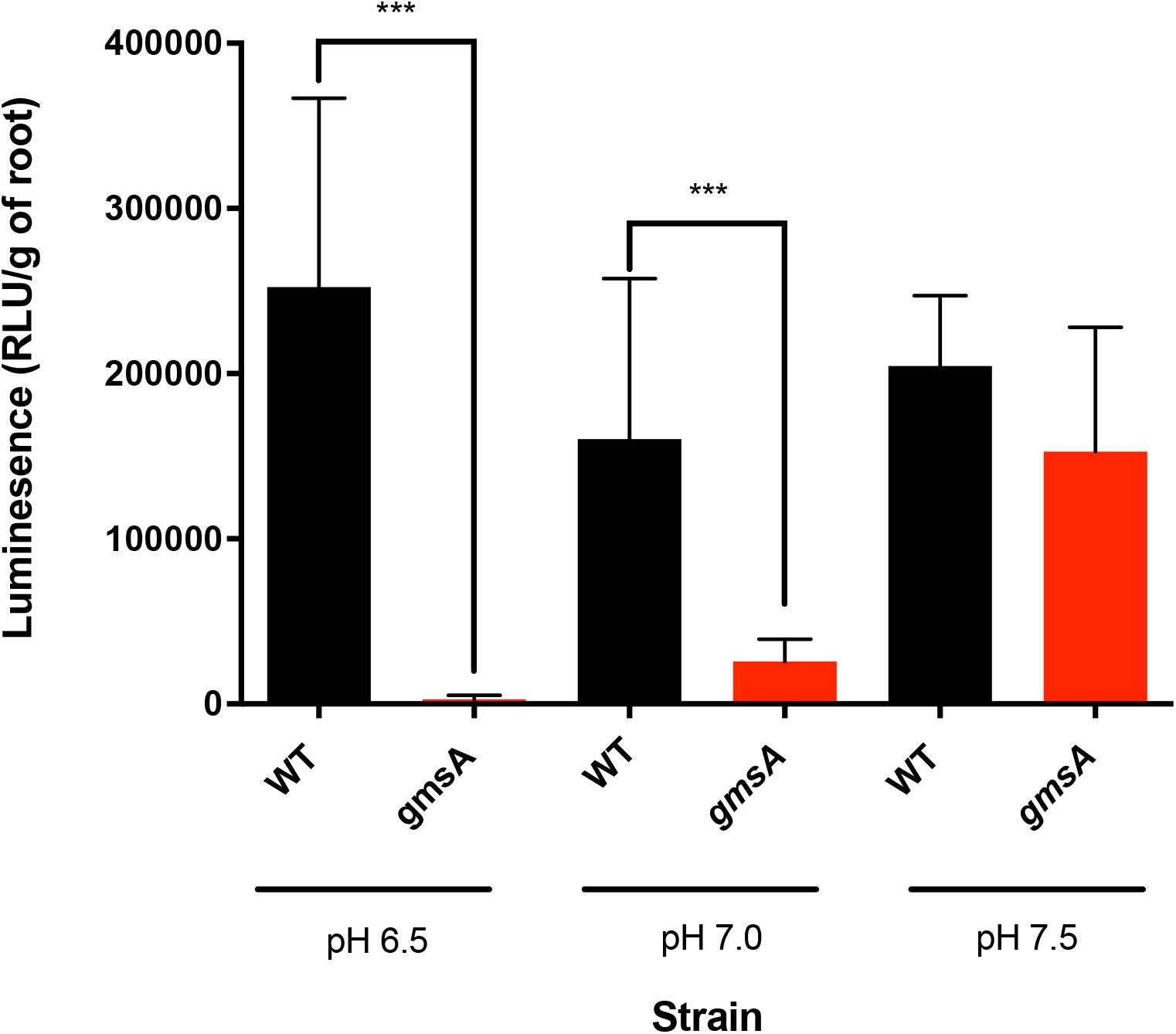
Effect of pH on primary attachment to pea roots. Attachment of Rlv3841 (WT) (black) and strain mutated in *gmsA* (RL1661) (red) at acidic, neutral and alkaline pH using the Lux-based whole root attachment assay. Luminescence (RLU/g of root) shows bacterial attachment after 1 h. *n* = 10. Data are expressed as the mean ± SD. ***= *p* < 0.001 using Student’s t-test. There is no statistically significant difference between primary attachment to roots of Rlv3841 (WT) at different pHs.

### *Mariner* transposon classification of genes affected in primary attachment

INSeq was used to investigate attachment of Rlv3841 to pea under acid (pH 6.5), neutral (pH 7.0) and alkaline (pH 7.5) conditions. The average input transposon insertion density (% of TA sites carrying one or more insertions) was 82%, while the averages for output densities were 48% (pH 6.5) and 50% (pH 7.0 and pH 7.5). Approximately 97% of the Rlv3841 genome (7,335 genes, encoded on a circular chromosome and six plasmids [23]), contains one or more TA sites available for INSeq analysis. In the input library 6,150 genes (87%) were classified neutral (NE). Genes in the input library classified as essential (ES) (< 0.1%), growth-defective (GD) (10%) and growth-advantaged (GA) (3%), were already compromised prior to attachment to plant roots and so not considered further.

For primary attachment at each pH (6.5, 7.0 and 7.5), 5,986 genes were assigned an INSeq classification: essential (ES), defective (DE), neutral (NE) or advantaged (AD) (Datasheet S1) through HMM analysis, using the same criteria defined for ES, GD, NE and GA, respectively [29] (the designations describing growth, e.g., growth defective (GD), were considered inappropriate for this study). The overall pattern of gene classification is remarkably similar at all pHs: ES 0.1-0.2%, DE 2.3-2.5%, NE 96-97% and AD < 0.01% (with approx. 1% failing to give a clear classification), although the genes falling into the categories at each pH is different. For genes lacking a consensus classification, manual curation was performed.

In total, 280 genes are required (classified ES/DE) for root attachment (Datasheet S2). Genes shown to be required for growth and survival in previous INSeq analyses [22, 47] were also removed, leaving 115 genes in the core “attach-ome” (Fig. 3, Supplementary Table S3). Twenty-two genes are required for attachment at pH 6.5, pH 7.0 and pH 7.5 (Fig. 3, Table 1), with genes required at specific pHs (Fig. 3) listed in Supplementary Tables S4-S9.

**Table 1.** Core attach-ome of 22 genes required for initial attachment to pea roots by Rlv3841 at pH 6.5, pH 7.0 and pH 7.5.

| Gene | Name | Description |
| --- | --- | --- |
| pRL100053* |  | Putative transmembrane protein with helix-turn-helix 37 domain |
| pRL100174 |  | Conserved hypothetical protein |
| RL0551 | *hslO* | Hsp33-like chaperonin HslO |
| RL0876 |  | Conserved hypothetical cytoplasmic protein |
| RL1381 |  | Uncharacterised protein. Unknown localization |
| RL1478 | *amn* | AMP nucleosidase; catalyses hydrolysis of AMP to form adenine and ribose 5-phosphate |
| RL2400 |  | Putative MarC family transmembrane protein, unknown function |
| RL2513 | *tpiA* | Putative triosephosphate isomerase TpiA |
| RL2637 | *recA* | DNA repair and SOS response RecA |
| RL3322 | *pfp* | Putative pyrophosphate-fructose 6-phosphate 1-phosphotransferase |
| RL3752* | *pssA* | Glycosyl transferase involved in EPS biosynthesis |
| RL3766 | *rpoH1* | RNA polymerase sigma-32-factor, heat shock sigma factor RpoH1 |
| RL3987 | *vapB* | SpoVT-AbrB domain-containing protein. Anti-toxin (VapB) of VapBC TA |
| RL3988 | *vapC* | PIN domain-containing protein. Single-stranded RNA nuclease. Toxin (VapC) of VapBC TA |
| RL3989 | *ruvA* | Holliday junction ATP-dependent DNA helicase RuvA |
| RL3990 | *ruvB* | Holliday junction ATP-dependent DNA helicase RuvB |
| RL4065 |  | Hypothetical cytoplasmic protein with no conserved domains |
| RL4145*<sup>a</sup> | *pckR* | LacI family transcriptional regulator |
| RL4362 |  | Putative cobalamin (vitamin B12) synthesis protein, CobW domain |
| RL4363 | *dacC* | Putative penicillin binding protein, peptidase S11 domain |
| RL4381 |  | Putative POTRA-domain transporter |
| RL4382* |  | Filamentous hemagglutinin adhesin |

\*Mutant made and strain assessed in Lux-based assay for initial attachment
<sup>a</sup> Gene encodes a regulator of gene transcription. RNASeq was performed on a strain mutated in this gene.

**Fig. 3.**
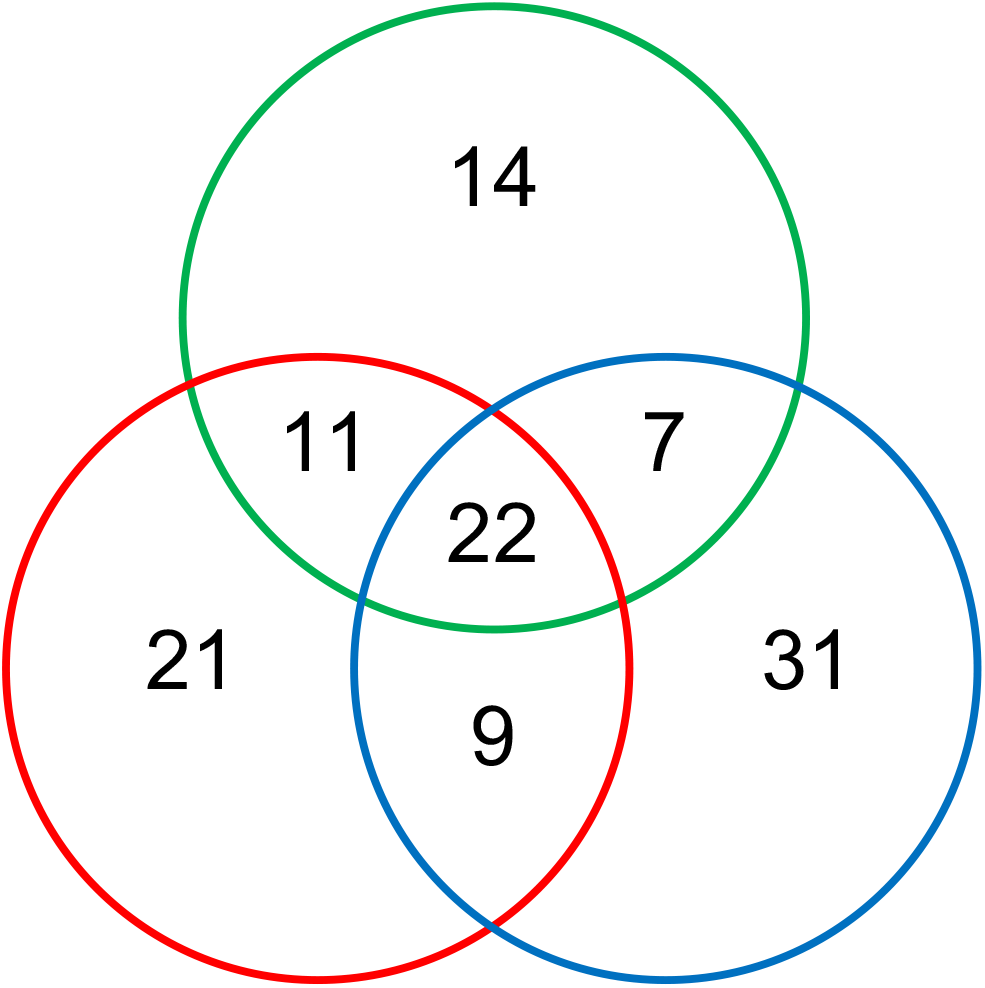
Rlv3841 genes required for primary attachment to pea roots under acidic, neutral and alkaline pH. A total of 115 genes were classified as required for attachment (ES/DE) in INSeq experiments performed at pH 6.5, pH 7.0 and pH 7.5 (Supplementary Table 3), with 22 genes being required for attachment at all pHs (Table 1). Colour of circle indicates pH; on the left-hand side (red) = pH 6.5, at the top (green) = pH 7.0 and on the right (blue) = pH 7.5. Genes are listed in Supplementary Tables S4-S9.

### Twenty-two genes required for attachment at all pHs make up the core attachome

Twenty-two genes are important under all conditions tested (Fig. 3) and can be considered as the core ‘attachome’ (their functions are summarised in Fig. 5). This core attachome (Table 1) contains membrane and the cell surface proteins, including the cell-surface filamentous hemagglutinin adhesin (RL4382), together with its transporter (RL4381), and MarC family transmembrane protein encoded by RL2400 (function unknown). Also required are enzymes involved in the biosynthesis of surface macromolecules, acidic EPS and capsular polysaccharide (RL3752 (*pssA*, glycosyl transferase)) and peptidoglycan (RL4363 (*dacC*)). Rhizobial mutants in *pssA* are deficient in acidic EPS production, form biofilms slowly compared to WT and do not attach to root hairs [4]. The biofilms that are formed by *pssA* mutants are flat and unstructured [7]. Enzymes affecting attachment include a triosephosphate isomerase and pyrophosphate-fructose 6-phosphate 1-phosphotransferase) (encoded by RL251 (*tpiA*) and RL3322 (*pfp*), respectively) (Table 1).

TpiA is upregulated in *Staphylococcus aureus* biofilm, possibly due to oxygen limitation [48]. Glycolytic enzymes can play additional roles when localised on the cell surface (e.g. α-enolase plasminogen binding in streptococci [49] and GAPDH transferrin binding activity in *S. aureus* [50]) and surface-localised TpiA has been shown to have a direct role in *Mycoplasma gallisepticum* attachment to host cells [51]. *R. leguminosarum* uses the Entner-Doudoroff pathway for sugar catabolism so while the reaction catalysed by Pfp is reversible, it is likely to be involved in gluconeogenesis converting D-fructose-1,6-diphosphate to D-fructose-6-phosphate. CobW domain-containing cobalamin synthesis protein is encoded by RL4362. In *E. meliloti*, biosynthesis of cobalamin is needed for symbiosis with *M. sativa* [52]. The proteins encoded by four other genes (pRL100174, RL0876, RL1381 and RL4076) remain uncharacterised (Table 1).

Many of the other genes in the core attachome (Table 1. Fig. 4) encode proteins involved in adaption to changing environments and managing resulting stress, e.g., RL0551 (*hslO*), a chaperonin whose function is to protect thermally unfolding and oxidised proteins from aggregation [53] and RL1478 (*amn*), an AMP nucleosidase which catalyses hydrolysis of AMP. Changes in AMP levels allow rapid adjustments to changing metabolic conditions [54]. Involved in DNA repair under changing environments are RL2637 (*recA*) (also responsible for the SOS response), with disruption of *recA* shown to reduce adherence and colonization of host cells by *Vibrio cholerae*, although the mechanism remains unknown [55], and RL3989 (*ruvA*) and RL3990 (*ruvB*) (RuvAB are Holliday junction ATP-dependent DNA helicases involved in DNA damage repair), shown to be involved the osmotic shock response [56]. Contiguous with RL3898-9 (*rvuAB*) (Table 1), are the overlapping open reading frames RL3987 (encoding a SpoVT-AbrB domain-containing protein of 87 amino acids) and RL3986 (encoding a PIN domain-containing protein of 133 amino acids), which have the characteristics of a toxin-antitoxin (TA) pair of the Virulence Associated Protein (VapBC) family [57]. Often sharing little primary sequence homology, other than three specific acidic residues, PIN domain-containing protein RL3988 (*vapC*) is the toxin, an RNase, cleaving single-stranded RNA in a sequence-specific, Mg^2+^- or Mn^2+^-dependent manner [57]. VapC is rendered inactive through protein-protein interactions with inhibitor RL3987 (*vapB*). A large number of VapBC TA systems are found in many procaryotes, including *Ensifer* (formerly *Sinorhizobium*) *meliloti* which has 21 such systems [57]. These include NtrPR, which has been shown to regulate a wide-range of metabolic transcripts, presumed to mediate symbiosis, and environmental and other stresses [58].

**Fig. 4.**
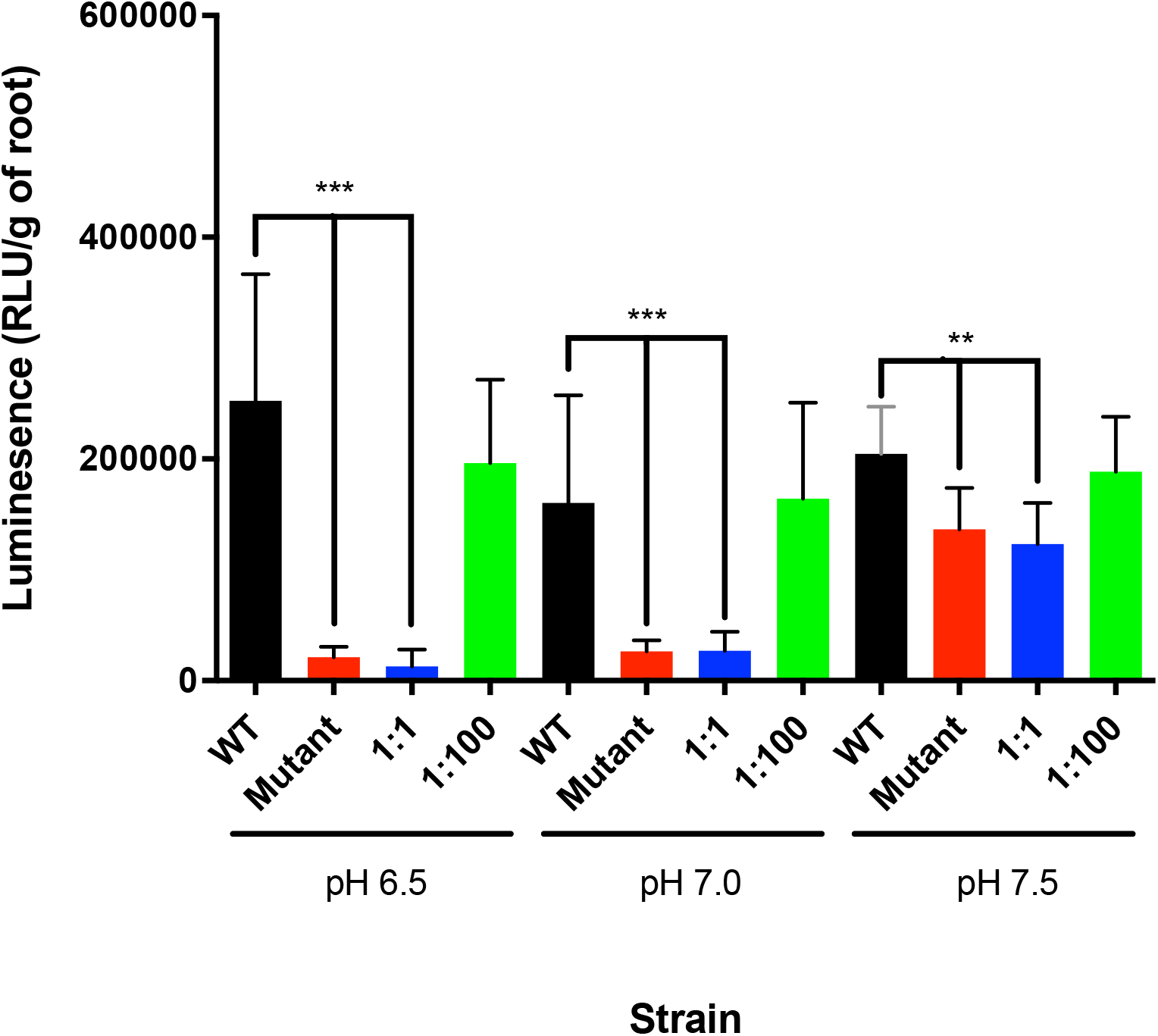
Effect of co-inoculation with wild-type bacteria on primary attachment to pea roots of a mutant at different pHs. Comparison of Rlv3841 (WT) (black) and a strain mutated in RL2969, as single inoculum (Mutant) (red), or in 1:1 (blue) or 1:100 (green) co-inoculation with unlabelled Rlv3841 using the Lux-based whole root attachment assay. Luminescence (RLU/g of root) shows bacterial attachment after 1 h. *n* = 10. Data is displayed as mean ± SD. ***= *p* < 0.001, **= *p <* 0.005 using Student’s t test. An unpaired t-test was used to compare groups.

Genes encoding proteins involved in controlling gene transcription, some potentially with a role in adaption and tolerance to hostile/stressful conditions, include the LacI family transcriptional regulator RL4145 (*pckR*), transmembrane protein pRL100053 (a membrane protein with a helix-turn-helix domain) and alternative sigma factor (heat-shock sigma factor) RL3766 (*rpoH1*) (Table 1). Rlv3841 has two genes encoding RpoH; RL3766 (*rpoH1)* and RL4614 (*rpoH2*) which show 47% amino acid identity to each other. This is similar to *E. meliloti*, where RpoH1 and RpoH2 share 42% identity [59], and 87% and 75% identity, respectively, with their Rlv3841 homologues. In *E. meliloti*, only RpoH1 complements mutation of the *E. coli* heat-shock sigma factor and only mutation of *rpoH1* leads to a *S. meliloti* Fix-phenotype [59]. RL4614 (*rpoH2*) was not required for attachment to roots (Datasheet S1). Sequence motifs identified for genes regulated by RpoH1 and RpoH1/H2 of *E. meliloti* [60] were used to probe the Rlv3841 genome (RpoH1 shown in Supplementary Table S10 and RpoH1/H2 in Supplementary Table S11). Five genes required for root attachment have RpoH1 recognition sequences upstream; pRL100053 (Tables 1 & 4), RL2520 (Table 4), RL2694 (Table 4) and RL3766 (*rpoH1*) (Tables 1 & 4) and RL4062 (Supplementary Tables S3 & S6). Interestingly, RpoH1 regulates its own expression. Genes pRL120322 (*fhuA2*) (Supplementary Tables S3 & S6) and RL4382 (Tables 1 & 4) have the motif for RpoH1/H2 binding. These genes are required at all pHs, with the exception of pRL120322 (*fhuA2*) and RL4062 (pH 7.5 only), RL2520 (pH 6.5 and pH 7.0) and RL2694 (pH 7.0 only). It is highly likely that the requirement for RpoH1 for root attachment in Rlv3841 is because its mutation disrupts this regulatory network, blocking expression of genes downstream.

### Genes required for root attachment vary with environmental pH

There are substantial differences in the gene requirements for attachment under different pH conditions (Fig. 3). One example is the requirement for glucomannan only under acid and neutral conditions, and not in an alkaline environment [4]. Our INSeq results for RL1661 (*gmsA*) of DE at pH 6.5 and pH 7.0, and NE at pH 7.5 (Table 2) are in complete agreement with the described phenotype [4]. Mutation of RL1661 (*gmsA*) encoding glucomannan biosynthesis protein A prevents bacteria from attaching to pea roots at pH 6.5 and pH 7.0 (Table 2).

**Table 2.**
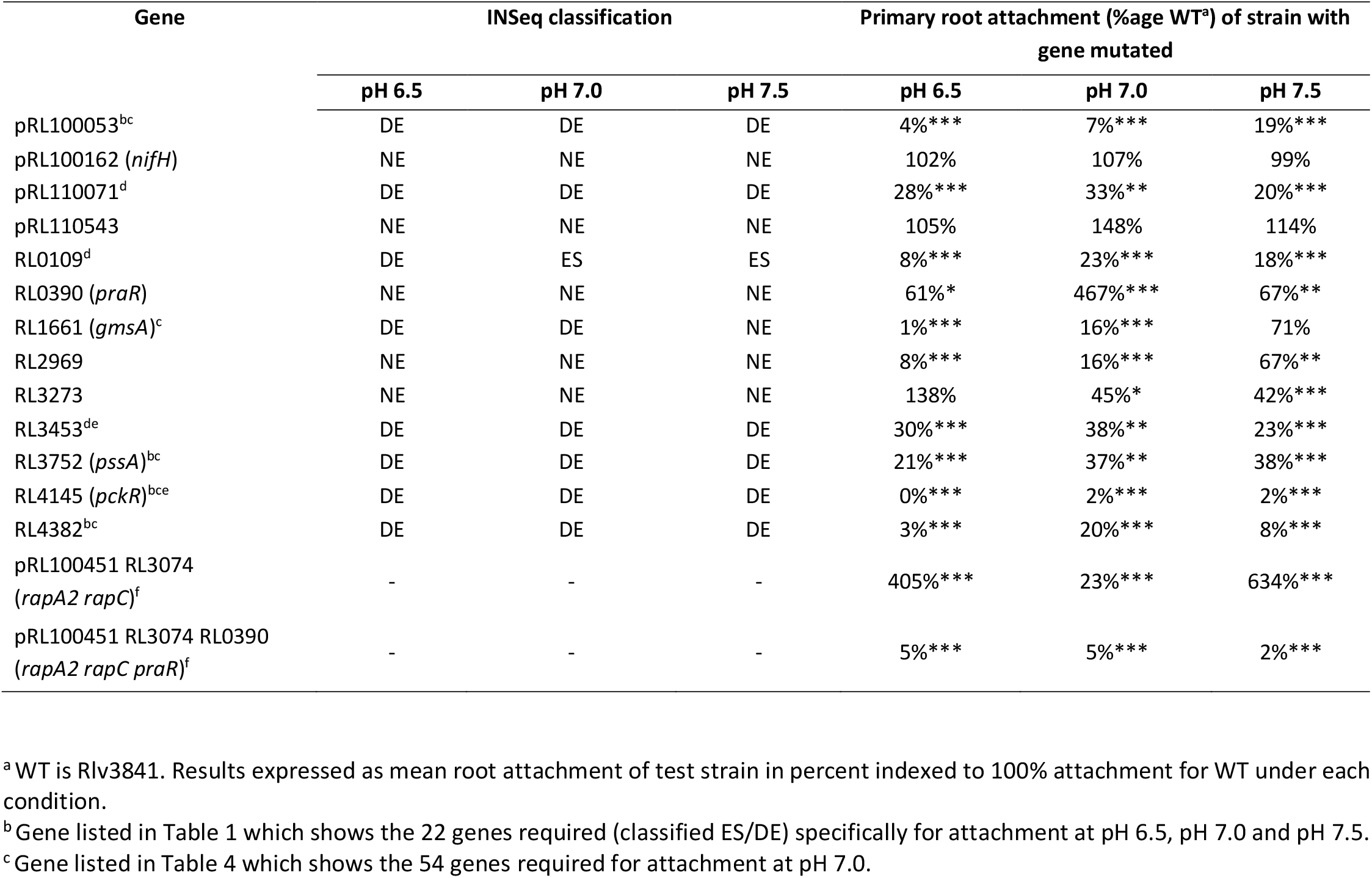
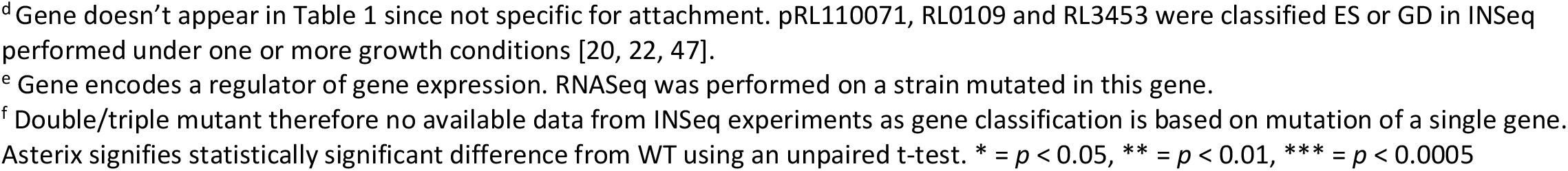
Comparison of INSeq classification of gene and primary attachment to roots of a strain with the gene mutated.

| Gene | INSeq classification |  |  | Primary root attachment (%age WT<sup>a</sup>) of strain with gene mutated |  |  |
| --- | --- | --- | --- | --- | --- | --- |
|  | pH 6.5 | pH 7.0 | pH 7.5 | pH 6.5 | pH 7.0 | pH 7.5 |
| pRL100053<sup>bc</sup> | DE | DE | DE | 4%*** | 7%*** | 19%*** |
| pRL100162 (*nifH*) | NE | NE | NE | 102% | 107% | 99% |
| pRL110071<sup>d</sup> | DE | DE | DE | 28%*** | 33%** | 20%*** |
| pRL110543 | NE | NE | NE | 105% | 148% | 114% |
| RL0109<sup>d</sup> | DE | ES | ES | 8%*** | 23%*** | 18%*** |
| RL0390 (*praR*) | NE | NE | NE | 61%* | 467%*** | 67%** |
| RL1661 (*gmsA*)<sup>c</sup> | DE | DE | NE | 1%*** | 16%*** | 71% |
| RL2969 | NE | NE | NE | 8%*** | 16%*** | 67%** |
| RL3273 | NE | NE | NE | 138% | 45%* | 42%*** |
| RL3453<sup>de</sup> | DE | DE | DE | 30%*** | 38%** | 23%*** |
| RL3752 (*pssA*)<sup>bc</sup> | DE | DE | DE | 21%*** | 37%** | 38%*** |
| RL4145 (*pckR*)<sup>bce</sup> | DE | DE | DE | 0%*** | 2%*** | 2%*** |
| RL4382<sup>bc</sup> | DE | DE | DE | 3%*** | 20%*** | 8%*** |
| pRL100451 RL3074<br>(*rapA2 rapC*)<sup>f</sup> | - | - | - | 405%*** | 23%*** | 634%*** |
| pRL100451 RL3074 RL0390<br>(*rapA2 rapC praR*)<sup>f</sup> | - | - | - | 5%*** | 5%*** | 2%*** |

<sup>a</sup> WT is Rlv3841. Results expressed as mean root attachment of test strain in percent indexed to 100% attachment for WT under each condition.
<sup>b</sup> Gene listed in Table 1 which shows the 22 genes required (classified ES/DE) specifically for attachment at pH 6.5, pH 7.0 and pH 7.5.
<sup>c</sup> Gene listed in Table 4 which shows the 54 genes required for attachment at pH 7.0.
<sup>d</sup> Gene doesn't appear in Table 1 since not specific for attachment. pRL110071, RL0109 and RL3453 were classified ES or GD in INSeq performed under one or more growth conditions [20, 22, 47]. <sup>e</sup> Gene encodes a regulator of gene expression. RNASeq was performed on a strain mutated in this gene. <sup>f</sup> Double/triple mutant therefore no available data from INSeq experiments as gene classification is based on mutation of a single gene. Asterix signifies statistically significant difference from WT using an unpaired t-test. \* = $p < 0.05$ , \*\* = $p < 0.01$ , \*\*\* = $p < 0.0005$

### Genes needed under specific conditions

There are 21 genes specifically required for root attachment only at pH 6.5 (Supplementary Table S4). Amongst these are RL0032 (*npr*) and RL0033 (*manX*) which encode part of the phosphenolpyruvate phosphotransferase system (PTS). Mutants in PTS genes have a ‘dry’ morphology due to reduced secretion of EPS [61]. The reduction in EPS in these mutants may be even greater at pH6.5 than at pH 7 or pH 7.5, leading to defective attachment. Likewise, pRL100274 (*fucA*), which encodes an α-L-fucosidase which cleaves fucosidic bonds in glycans (particularly in peptidoglycan structures) [62] and is likely involved in remodelling of the cell surface. Furthermore, there is a requirement for RL0398, encoding a putative N-acetyltransferase, which may be involved in peptidoglycan modification, RL0726 (putative transglycosylase) able to degrade peptidoglycan via β 1-4 glycosidic bond cleavage [63], RL3179, encoding a putative cobalamin (vitamin B12) synthesis protein (which may be involved in peptidoglycan amidation) and RL4404 (*gelA*) encoding gel-forming EPS production protein [4]. A requirement for these genes suggests the importance at pH 6.5 for the presence of cell surface peptidoglycan, as well as modification and remodelling. At this pH, plant lectins are present on roots, while at alkali pH they dissociate [6]. Membrane proteins are also needed at pH 6.5, e.g., RL2780 (*dgkA*) (DgkA is involved in the turnover of membrane phospholipid, and known to be important for gram negative bacterial adhesion to host cells [64]) and pRL120475 (*impA*), an inner membrane protein. Also, RL2316, a putative guanylate cyclase which catalyses formation of cyclic di-GMP (c-di-GMP), a key secondary messenger in biofilm formation or motile to sessile lifestyle switch [65]. Biofilm formation is regulated by c-di-GMP levels in *Shewanella oneidensis* [66].

Fourteen genes are required specifically at pH 7.0, including membrane protein RL1371 (Supplementary Table S5).

Thirty-one genes are needed for root attachment specifically at pH 7.5 (Supplementary Table S6). As with genes required for attachment at pH 6.5, the importance of the bacterial cell surface is apparent. There is a requirement for genes encoding proteins involved in peptidoglycan metabolism e.g., RL2489A, encoding a transglycosylase-associated protein, and RL2778, encoding an exoplysaccharide biosynthesis protein. Also, RL2644 (*sixA*, the only known bacterial phosphohistidine dephosphorylase), part of the Npr system which acts to dephosphorylate Npr in *E. coli* [67] is implicated in biofilm formation [68].

Interestingly, Npr itself is required for attachment at pH 6.5 (Supplementary Table S4), while at pH 7.5, lack of SixA, which will leave Npr phosphorylated, is detrimental to bacterial attachment. One possibility is that phosphorylated Npr inhibits attachment at pH 7.5 but promotes it at pH 6.5. The composition of membranes is important, RL1338 (*pmtA*), a phosphatidylethanolamine N-methyltransferase involved in membrane lipid phosphatidylcholine synthesis being important at pH 7.5. A role in attachment is also suggested by its expression being significantly upregulated in a *praR* mutant of Rlv3841 [12]. In *Bradyrhizobium japonicum*, mutation of *pmtA* disrupts symbiosis [69]. RL4018 encodes an ATP-binding component of an ABC transporter showing 96% identity to lipid A exporter from *R. leguminosarum* biovar *trifolii* WSM2304. Mutants with reduced lipid A are delayed in nodulation and impaired in bacteroid shape [70]. Furthermore, in Rlv3841, defects in lipid A modification have reduced surface attachment and motility, although no effect on symbiosis was observed [71]. Interestingly, genes RL2975-7 encoding an ABC transporter showing reduced EPS export and found to be important maintaining protection against desiccation [72], are classified NE for attachment at all pHs examined here (Datasheet S1).

Outer membrane siderophore receptor FhuA2 (pRL120322) is required at pH 7.5 only, but the significance of this is unclear. RL3253 (*hflC*) and RL3254 (*hflK*) encode transmembrane proteases which modulate activity of HflB. HflB (or FtsH) is an AAA metalloprotease, involved in membrane protein regulation, LPS biosynthesis and biofilm formation in *E. coli, B. subtilis* and other bacteria [73, 74], with a role in membrane regulation and biofilm formation [75, 76]. HflB (encoded by RL3965) is ES/DE under all conditions, including the input library. As for control of transcription, RL1505 (*rpoZ*) encoding the smallest subunit (omega) of DNA-directed RNA polymerase, is specifically required only at pH 7.5. It has been shown that lack of RpoZ impairs biofilm formation and sliding motility in *Mycobacterium smegmatis* [77] and biofilm formation in *E. coli* (via ppGpp and the stringent response) [78] and also in Gram-positive *S. aureus* through transcriptional changes probably due to the lack of stability of the resulting omega-lacking RNA polymerase complex [79].

Genes required at pH 6.5 & pH 7.0 include RL1661 (*gmsA*), well-documented to be required only for attachment at acid and neutral pH [4], RL1600 (*ppx*) an exopolyphosphatase, putative transport proteins pRL110043 and RL2520, and membrane proteins RL3277 and RL4309 (Supplementary Table S7). Genes required at pH 7.0 & pH 7.5 or at pH 6.5 & pH 7.5 are listed in Supplementary Table S8 and S9, respectively.

### Importance of genes required for attachment in other steps of symbiosis

Comparison of all 54 genes required for attachment at pH 7.0 with the data of Wheatley et al. [23], shows many are crucial at other stages of rhizobial symbiosis (Table 3, Fig. 5). While many of the genes in the core attachome are specific for primary attachment and not required at any other stages of the symbiosis process (pRL100174, RL0551 (*hslO*), RL0876, RL1381, RL2513 (*tpiA*), RL3752 (*pssA*), RL4145 (*pckR*) and RL4381-2), several are involved at one or more other stages of symbiosis (RL2400, RL2637 (*recA*), RL3322 (*pfp*), RL3766 (*rpoH1*), RL4065 and RL4362-3 (*dacC*)) or throughout the whole development process (pRL100053, RL1478 (*amn*) and RL3987-90) (Table 3).

**Table 3.**
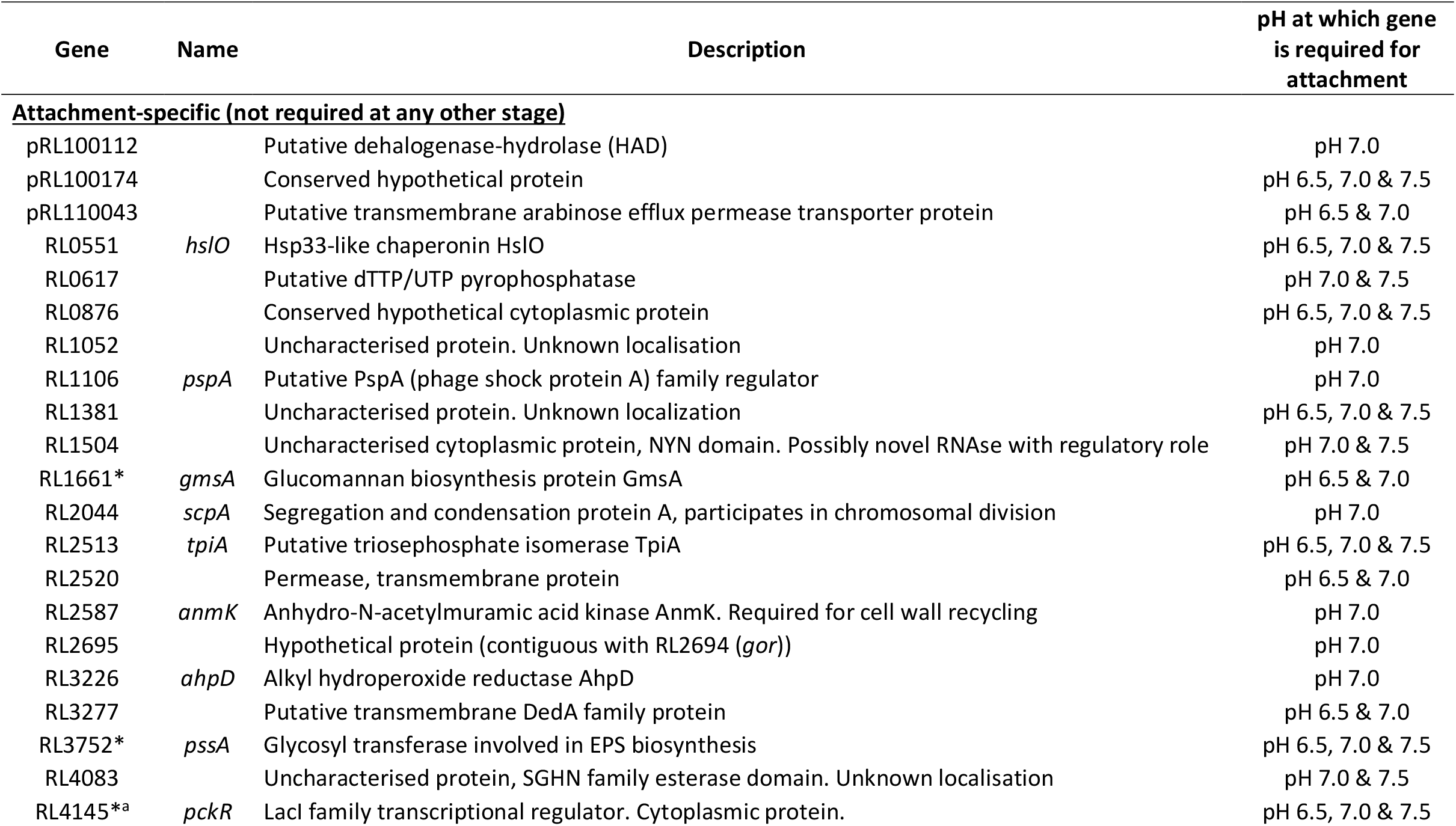
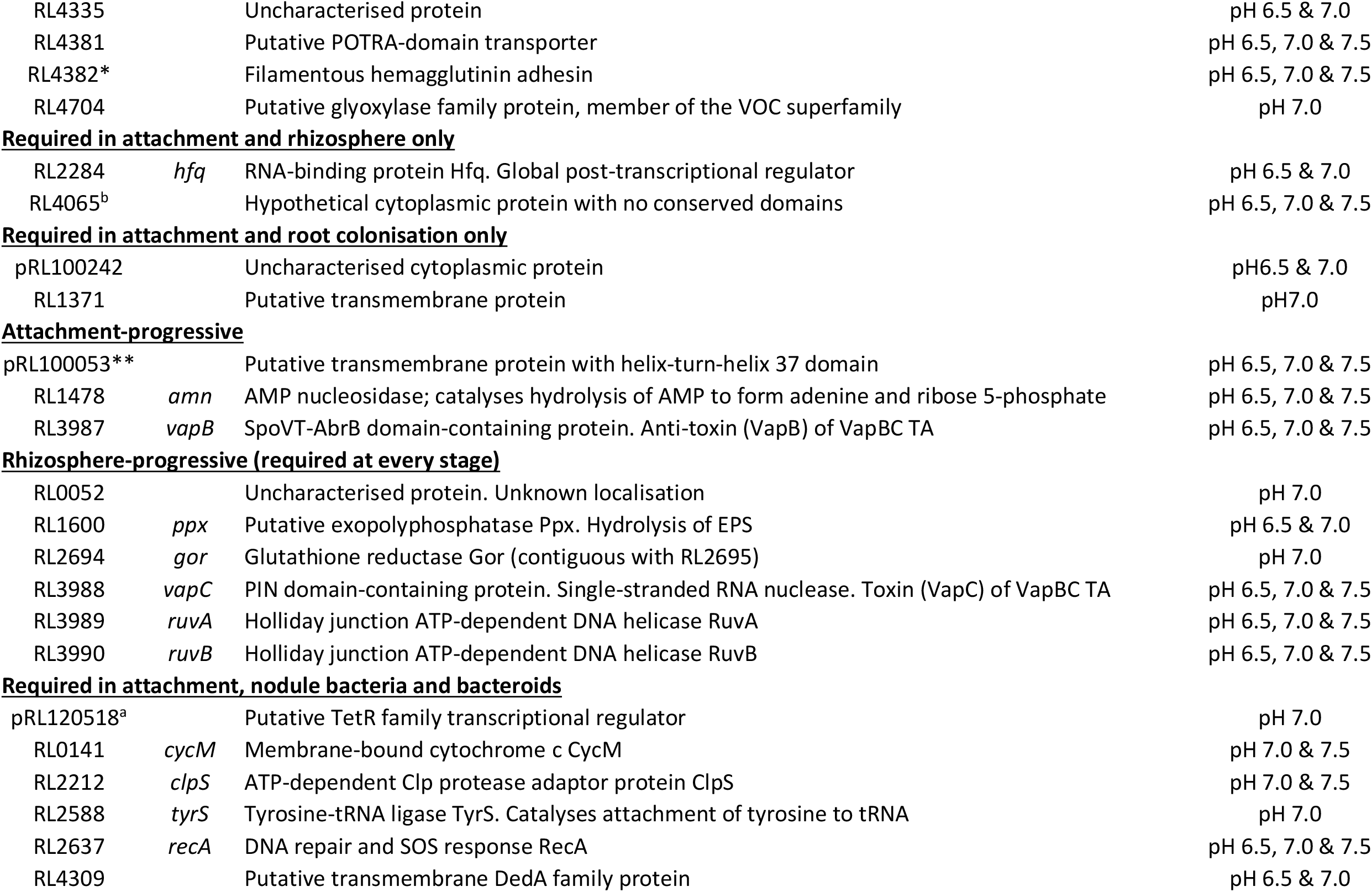
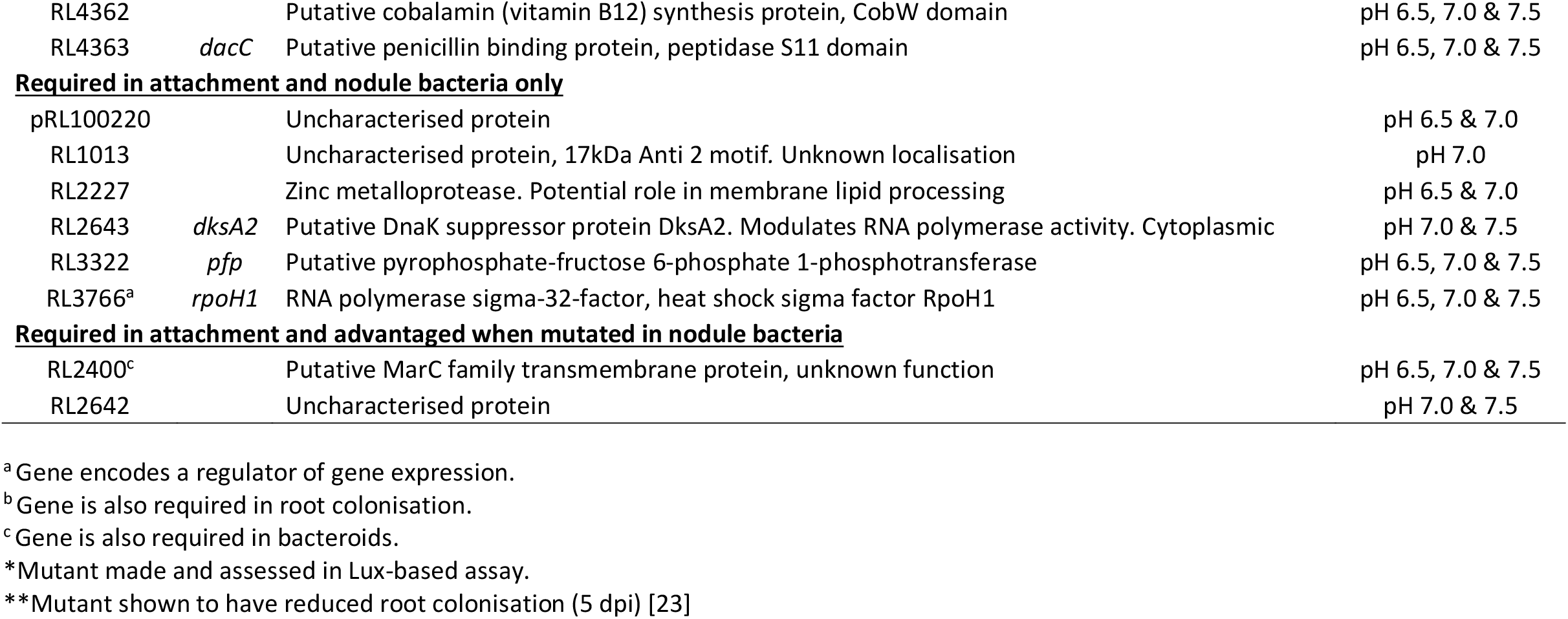
Genes required for root attachment at pH 7.0 grouped by their requirement at different stages (rhizosphere, root colonisation, nodule bacteria and bacteroids) of nodule development.

**Fig. 5.**
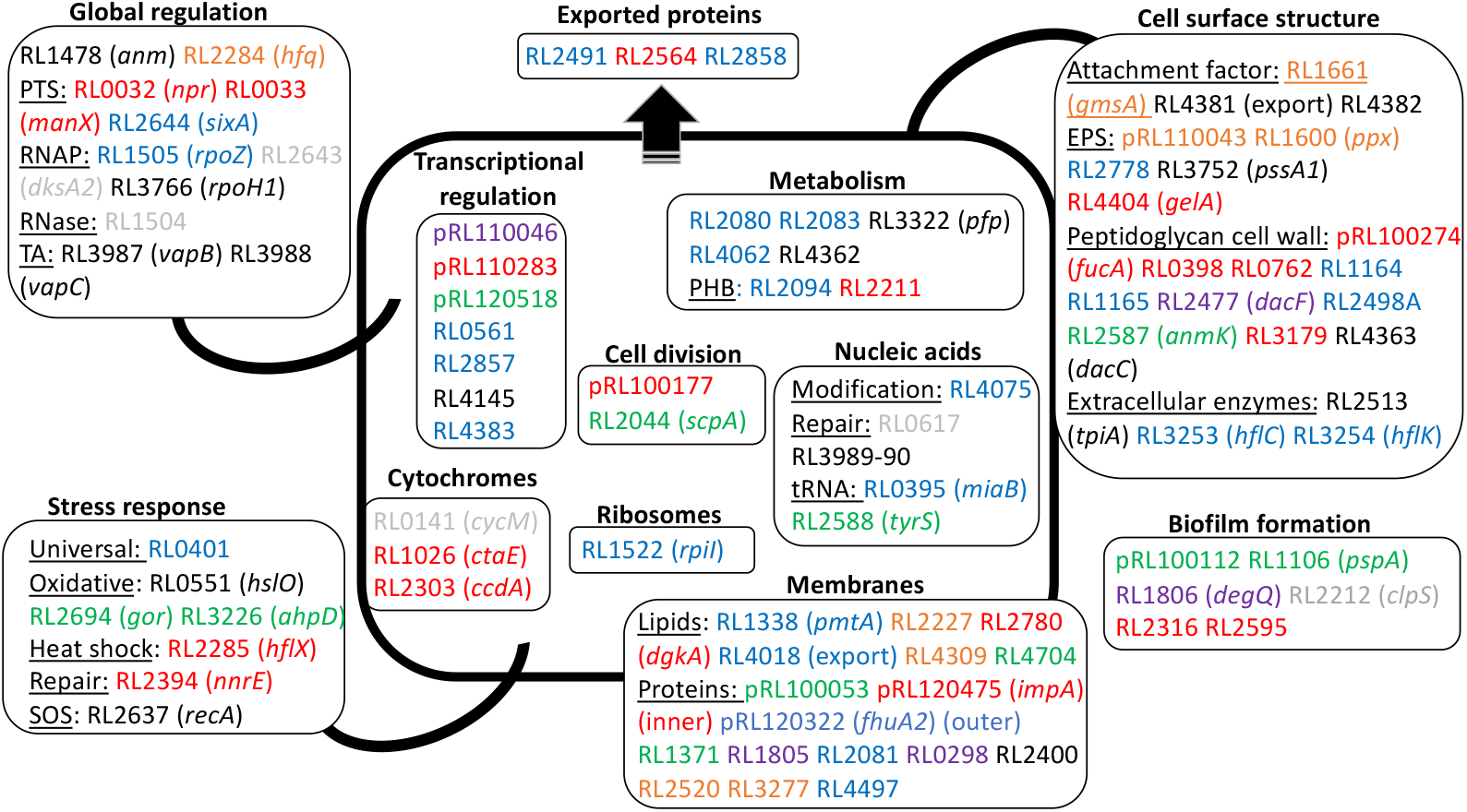
Summary of Rlv3841 genes required for initial attachment to pea roots at acidic, neutral and alkaline pH grouped by function. From INSeq, the 115 genes required at pH 6.5 to pH 7.5 (Fig. 3, Supplementary Table S3), are shown grouped by their involvement in different cellular structures or processes, and colour coded: pH 6.5 (red), pH 7.0 (green), pH 7.5 (blue), at pH 6.5 and pH 7.0 (orange), pH 7.0 and pH 7.5 (grey) and pH 6.5 and pH 7.5 (purple). In addition, there are 24 genes which have an, as yet, unidentified function (Supplementary Table S3).

As the rhizosphere to symbiosis INSeq experiments were performed under neutral conditions [23], only attachment at pH 7.0 was considered. Almost half of the 54 genes required for initial attachment at pH 7.0 fall into the category of being required for initial attachment, but having no further effect on the development of nitrogen-fixing bacteroids (Table 3, Fig. 6). These include RL1661 (*gmsA*), which is classified as NE throughout nodule development [23] (Table 3), and illustrates that while RL1661 (*gmsA*) plays a role in initial attachment at pH 7.0 it does not alter nodulation competitiveness which proceeds via root hair attachment.

**Fig. 6.**
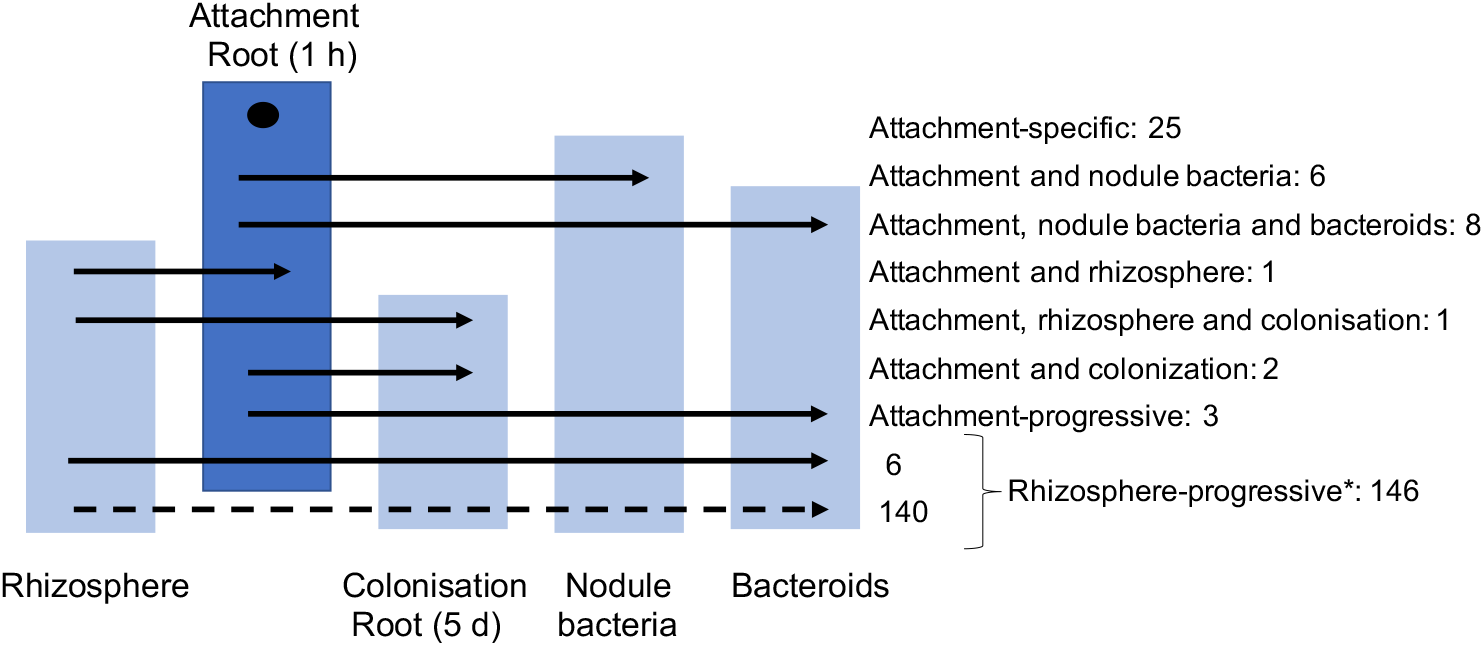
Involvement of genes required for root attachment at pH 7.0 at other stages of nodule development (in the rhizosphere, for root colonisation, in nodule bacteria and in bacteroids). Fifty-four genes are required for initial root attachment (Fig. 3) and their involvement at other stages of infection and symbiosis was determined from the INSeq study of nodule development [23]. Twenty-five genes are root-attachment specific, with the remainder required at one or more stages of symbiosis (attachment to roots (1 h) is represented by dark blue box, while those of other developmental stages are pale blue). Two genes (RL2400 and RL2642) are not represented; they are INSeq-classified as AD in nodule bacteria [23] (Table 3). *Rhizosphere-progressive genes, in total 146 [23], with six of them also required for initial attachment.

Some attachment genes are also required in the rhizosphere and/or colonisation of pea roots (after 5 d). The global post-transcriptional RNA-binding protein Hfq encoded by RL2284 (*hfq*) is required for bacterial survival in the rhizosphere as well for attachment, but at no other stages of nodule formation, including root colonisation (Table 3). In *E. meliloti*, deletion of *hfq* delays nodulation and reduces competitiveness for attachment to alfalfa roots [80].

Employing the terminology used by Wheatley et al. [23], there are groups of genes that are ‘progressive’ in the process of nodule development. We can consider that the primary attachment described here forms a stage between those of the rhizosphere and 5-d root colonisation (Fig. 6). Gene pRL100053 was classified as colonisation-progressive and a single mutant was unable to colonise roots [23]. Here we show it is, in fact, affected earlier in the developmental process. It should be considered as attachment-progressive as shown by both INSeq designation and Lux assay with only 4-19% attachment of that of WT bacteria (Table 2). Others in this same category of attachment-progressive genes are RL1478 (*amn*) (discussed above) and RL3987 (uncharacterised protein) (Table 3).

Six bacterial genes required for attachment (RL0052, RL1600 (*ppx*), RL2694 (*gor*) and RL3988-90) were shown to be required for nodulation (Table 3, Fig. 6), falling into the group described as rhizosphere-progressive [23]. Of those with known functions, the three gene cluster RL3988, RL3989-90 (*ruvAB*) encodes proteins concerned with DNA repair associated with bacterial stress (discussed above), while RL0052 (*ppx*) encodes an exopolyphosphatase and may be involved in hydrolysis of EPS. RL1600 (*gor*) encodes a glutathione reductase, likely to be playing a role in cellular control of reactive oxygen species (Table 3).

Two further groups of genes required for attachment at pH 7.0 are also needed at other stages of the nodulation process, either in both bacteroids and nodule bacteria (8 genes) or only in nodule bacteria (6 genes) (Table 3). While it is clear that initial root attachment is not the same as attachment to root hairs, root attachment may be important for bacterial competition and long-term survival in the soil.

As the incubation time on roots was only 1 h, most of the responses in this study will not involve root induction of attachment factors. For example, even in an optimised temperature-regulated response system in *E. coli*, marked gene induction was not seen in under two hours [81]. Expression of genes required for attachment is either constitutive or occurs in the bacterial growth conditions prior to the assay (Fig. 3). As only approx. 17% of the genes required for initial attachment to roots are plasmid-encoded, despite making up approx. 35% of the Rlv3841 genome, there is a bias towards chromosomal genes in primary attachment of Rlv3841 to pea roots. However, plasmid genes, particularly of pRL8, are upregulated in the pea rhizosphere with 24 h post-inoculation [13] so plasmid genes may be involved in later stages (and possibly host-specific) of secondary attachment and colonisation of host roots, although longer time-scale (>24 h) INSeq secondary attachment/ colonisation studies would be needed to address this.

### Assessment of INSeq classification of genes using the Lux-based attachment assay

For a selection of genes, comparison was made between their INSeq classification and the effect of their mutation on bacterial root attachment. Strains were made by mutating individual INSEq-identified ES/DE genes pRL100053, pRL110071, pRL110543, RL0109, RL2969, RL3273, RL3453, RL4145 (*pckR*) and RL4382, which were assessed for root attachment, together with strains mutated in pRL100162 (*nifH*) [25], RL0390 (*praR*) or RL3752 (*pssA*) (Table 2). Under every pH condition there is excellent consensus between INSeq-classification and root attachment for pRL100153, pRL110071, RL0109, RL3752 (*pssA*), RL3453, RL4145 (*pckR*) and RL4382, which show reduced primary root attachment at pH 6.5, pH 7.0 and pH 7.5 (Table 2). In addition, there is agreement in the INSeq classification of RL1661 (*gmsA*) (DE at pH 6.5 and pH 7.0, NE at pH 7.5) and root attachment of the mutant strain (Table 2). RL0390 (*praR*) is classified NE but its mutation affects primary attachment (Table 2). However, as six TA sites is the minimum for assignment in HMM analysis [82, 83] and RL0390 (*praR*) only contains four TA sites, any predictions made from the INSeq results are not valid for this gene.

Furthermore, while Lux-based assays have only a single mutant strain the INSeq library has approximately 115,000 (82% insertion density of 140,845 genomic TA sites) mutant strains competing to attach to roots. Some mutants will be complemented *in trans* by diffusible factors, or aided in biofilm formation (e.g., by a surface polysaccharide). To test if differences in experimental setup has given an apparent discrepancy in results, a Lux-labelled mutant (RL2969 or RL3273 (Table 2)) strain was assessed for its ability to attach to roots in the presence of unlabelled WT bacteria (at ratios of WT:mutant of 1:1 and 100:1). The strain mutated in RL2969 regained the ability to attach to roots in the presence of a 100-fold excess of WT (Fig. 4). RL2969 encodes a putative transmembrane protein, all or part of which is perhaps released from the surface of WT cells. Alternatively, it might aggregate WT and mutant bacteria on the root surface, thereby complementing the mutant’s attachment-deficient phenotype. The strain mutated in RL3273 was not rescued by a 100-fold excess of WT bacteria, however this ratio is still much less than the 115,000-fold excess in INSeq experiments.

### Role of regulators in initial root attachment

Across the total range examined (pH 6.5-pH 7.5), there is a requirement at one or more condition for seven transcriptional regulators; pRL110046 (FNR/CRP family), pRL110283 (ArsR family), pRL120518 (TetR family), RL0561 (AraC family), RL2857 (LysR family), RL4145 (*pckR*) (LacI family), RL4383 (AsnC family) (Supplementary Table S3).

RL4145 (*pckR*) was investigated by RNASeq on a mutant strain (see below). Given gene proximity and common INSeq profiles, we can speculate that pRL110046, required at pH 6.5 and pH 7.5, may control expression of contiguous pRL110045 (encoding an uncharacterised protein) (Supplementary Table S9). TetR regulator pRL120518 may control the contiguous genes pRL120516-14 (*qatX3W3V3*), encoding the glycine betaine Quaternary Amine Transporter (QAT) family ABC transporter [84]. While INSeq classification of genes pRL120516-14 (*qatX3W3V3*) are all NE there are five other *qat* operons in Rlv3841 which could build in ‘redundancy’ and allow subunits to replace one another to form a functional transport system. AsnC family regulator RL4383 may control expression of contiguous genes RL4382-1, encoding the filamentous hemagglutinin adhesin and its transporter, required for attachment at all pHs (Tables 1, 2 & 4). From a brief examination, regulators encoded by pRL110283, RL0561 and RL2857 and have no obvious target gene(s).

**Table 4.** Potential rhicadhesin-encoding genes determined from INSeq analysis or proteomic screening.

| Gene | Description | Predicted protein size (kDa) | INSeq classification |  |  | Ca<sub>2</sub><sup>+</sup>-binding domain<sup>a</sup> |
| --- | --- | --- | --- | --- | --- | --- |
|  |  |  | pH 6.5 | pH 7.0 | pH 7.5 |  |
| **<u>Proteomic analysis of crude adhesin<sup>b</sup></u>** |  |  |  |  |  |  |
| RL0770 | putative phasin, phasin-2 superfamily | 16 | NE | NE | NE | Yes |
| **RL1635\*** | outer membrane protein | 19 | DE | DE | DE | No |
| RL3968 (*pal*)* | OmpA family peptidoglycan associated lipoprotein | 19 | ES | DE | ES | No |
| **RL4441 (*omp19*)\*** | outer membrane protein Omp19 | 19 | DE | DE | DE | No |
| **RL4733\*** | conserved hypothetical protein | 17 | ES | DE | ES | Yes |
| **<u>INSeq analysis<sup>c</sup></u>** |  |  |  |  |  |  |
| RL1165 | conserved hypothetical protein | 12 | NE | NE | DE | Yes |
| RL1166 | putative ribonuclease-L-PSP family protein | 14 | NE | NE | DE | Yes |
| RL1339 | conserved hypothetical protein | 15 | NE | NE | DE | No |
| RL2095 | conserved hypothetical protein | 15 | NE | NE | DE | Yes |
| RL2643 (*dksA2*) | putative DnaK suppressor protein | 15 | NE | NE | DE | No |
| RL2858 | conserved hypothetical exported protein | 13 | NE | NE | DE | No |
| RL4383 | putative AsnC family transcriptional regulator | 16 | NE | DE | DE | Yes |

<sup>a</sup>Ca<sup>2+</sup>-binding activity predicted using the IonCom tool [31].
<sup>b</sup>Candidate genes were identified by LC-MS in a proteomic screen of proteins in the crude adhesin band (Fig. 7A).
<sup>c</sup>Genes identified as encoding candidate rhicadhesin(s) on the basis of their size (encoding a 12-16 kDa protein) and INSeq classification: NE under acid, NE/DE/ES under neutral and DE/ES under alkaline conditions.
\*Also required for growth in input library and other media and conditions [20, 22, 23, 47]. Shown in bold are genes for which mutation was attempted.

Two regulators, RL3453 (histidine kinase of a two-component sensor/regulator) and RL4145 (*pckR*, LacI family transcriptional regulator), classified by INSeq as required for initial root attachment at all pHs (although mutation of RL3453 also reduced growth on succinate at 21% oxygen [22]) (Table 1) were chosen for RNASeq to investigate their target genes. Their role has also been confirmed by the whole root attachment assay (Table 2). Strains OPS1907 (mutation in RL3453) and OPS1908 (mutation in RL4145 (*pckR*)) were investigated using RNASeq, together with strain RU4062 (pRL10062 *(nifH*) mutant, unaffected for attachment) (Table 2). Many genes are differentially regulated, especially in OPS1908 (123 up-and 599 genes down-regulated ≥ 5-fold), while OPS1907 has five genes up- and 370 genes down-regulated ≥ 5-fold (Supplementary Table S12).

In OPS1908 (RL4145 (*pckR*) mutant), the most highly differentially expressed genes (56-to 67-fold) are *zwf1-pgl-edd* (RL0753-51), encoding glucose-6-phosphate dehydrogenase, 6-phosphogluconolactonase and phosphogluconate dehydratase, respectively, enzymes of the Entner-Doudoroff pathway, used in *Rhizobiaceae* for synthesis of glucose. RL4145 (*pckR*) shows 89% identity to *pckR* (*smc0297*) of *E. meliloti* which has been shown to be a key regulator of central carbon metabolism [85]. In *E. meliloti*, PckR acts as a repressor of the Entner-Doudoroff pathway (*zwf-pgl-edd*) and an inducer of *pckA, fbaB* and *mgsA*, genes involved in gluconeogenesis [85]. In addition to differential regulation of f *zwf1-pgl-edd* (RL0753-51, 56-to 67-fold) the *R. leguminosarum pckR* mutant shows up-regulation of *eda2* (RL4162, approx. 7-fold), with concomitant down-regulation of *pckA* (RL0037, approx. 50-fold down) and *fbaB* (RL4012, approx. 20-fold down), which mirrors *E. meliloti [85]* (Supplementary Table S12). However, expression of *mgsA* (RL0183, approx. 2.8-fold up) contrasts with *E. meliloti* where PckR acts as a positive regulator [85]. Many other genes up-regulated >5-fold in this mutant encode integral and outer membrane proteins, enzymes performing polysaccharide polymerisation and modification (e.g., glucosyl-, galactosyl-transferases, transglycosylases), membrane transport systems (both uptake and export) and, notably, six clusters of genes encoding uptake systems for iron, including those encoding siderophore vicibactin synthesis and transport proteins (pRL120312-21), siderophore interacting protein (RL2916), haem synthesis and transport (RL3692-7) and components of four ABC transporters of the FeCT family; pRL120711-2, pRL100326-7 (*fhuBD*), RL3698-70 (*hmuTV*) and RL2713-5 (Supplementary Table S12). Amongst the more than five hundred genes >5-fold down-regulated in OPS1908, which include numerous membrane proteins and transport systems, RL0933 (transmembrane protein) and RL0469 (exported surface protein) are the most down-regulated (approx. 300-fold and approx. 50-fold, respectively). It seems clear that unregulated carbon metabolism resulting from mutation of the key regulator PckR leads to widespread changes affecting both the cell surface and compounds transported across it. It seems that rather than controlling expression of any specific attachment factor, mutation of RL4145 (*pckR*) affects the bacterial surface, leading to the observed phenotype (Table 2).

Examining the expression in strain OPS1908 of genes identified as affecting attachment to roots shows RL3322 (*pfp*) encoding pyrophosphate--fructose 6-phosphate 1-phosphotransferase, an enzyme involved in carbon metabolism is unaffected by mutation of RL4145 (*pckR*), while expression of RL3752 (*pssA*), involved in exopolysaccharide biosynthesis, is up-regulated (approx. 5-fold) (both are required at all pHs, Table 1).

In strain OPS1907 (RL3453 mutant), vicibactin synthesis proteins encoded by pRL120315-6 are up-regulated approx.6-fold (also up-regulated in OPS1908, approx. 20-fold). Unique to OPS1907, is up-regulation of RL3181 encoding an RcnB family membrane protein (approx. 6-fold) and pRL100469 encoding a IS30 transposase (approx. 5-fold). As in OPS1908, the gene encoding transmembrane protein RL0933 is the most highly down-regulated gene (approx. 250-fold), with 339 of the 370 down-regulated genes in OPS1907, over 90%, also down-regulated (≥ 5-fold) in OPS1908 (Supplementary Table S12). These results show a huge overlap in the effects of these two different mutations in genes encoding regulators of gene expression, which result in a loss of ability to attach to roots, likely to be as a result of radical changes to the bacterial cell surface.

### Investigation of the gene encoding rhicadhesin

In an attempt to identify the gene encoding rhicadhesin [30], we performed cell fractionation and isolation of protein fractions as previously described in the original work [30] (Fig. 7A). The crude adhesin fraction isolated from the protein gel by excising the band running at approx.14 kDa (Fig. 7A). Following pre-incubation of pea root sections with the crude adhesin protein fraction, attachment of WT Rlv3841 was significantly reduced (Fig. 7B).

**Fig. 7.**
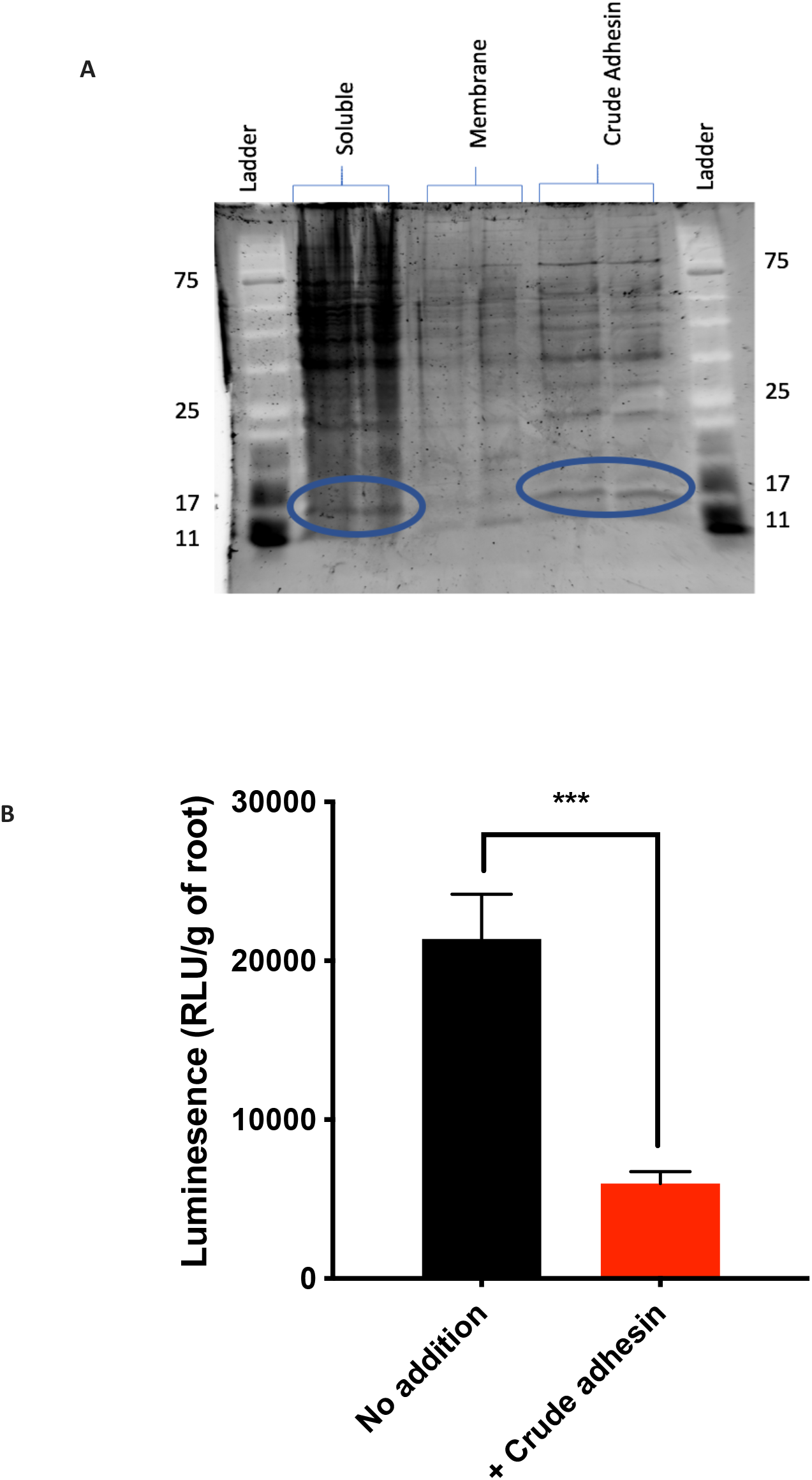
Purification of a crude adhesin protein fraction and its effect on primary attachment of Rlv3841 (WT) to pea roots. **A** SDS-gel of Rlv3841 cell fractions (soluble, membrane and crude adhesin) stained with SYPRO Ruby. The putative rhicadhesin fraction, a protein band at approx. 14 kDa (circled in blue), is visible in both soluble and crude adhesin fractions, the latter of which was excised from the gel and used for proteomic analysis and pre-incubation of roots. Ladder protein sizes (kDa) are indicated. **B** Root attachment of Rlv3841 (WT) using the Lux-based whole root attachment assay, with no addition (black) and pre-incubation of roots with crude adhesin for 1 h (+ Crude adhesin, red). Luminescence (RLU/g of root) shows bacterial attachment after 1 h. Data is displayed as mean ± SD, *n* = 5. ***= *p* < 0.001 using Student’s t test.

Analysis by LC-MS of the crude adhesin fraction isolated contained fifteen proteins (Supplementary Table S13). The five most likely candidates for rhicadhesin (sizes range from 16-19 kDa) are shown in Table 4. RL4733 was the most abundant protein, with RL3968 (*pal*) (approx. 50% of RL4733) and RL0770, RL1635 and RL4441 (*omp19*) (each approx. 10% of RL4733) (Supplementary Table S13). None of these has the expected attachment pH profile for the rhicadhesin gene; INSeq classification of RL0770 was NE for attachment at all pHs, while RL4733, RL3968 (*pal*), RL1635 and RL4441 (*omp19*) were classified ES/DE under all attachment conditions tested (Table 4). In addition, these latter four genes are also non-NE in both the input library and for growth on TY and other media [20, 22, 23, 47], indicating that they are not specifically involved in root attachment.

## Discussion

Bacteria attach initially to plant roots using mechanisms that are heavily dependent on environmental pH. We have been able to ascertain how disruption of multiple primary attachment factors affect downstream events leading to symbiosis. It is apparent there are layers of control of gene expression, both specific and global, which would benefit from other approaches to investigate the underpinning gene networks in detail. This research has initiated a deeper understanding of how environmental pH affects rhizobial attachment to roots, a key interaction between bacteria and plants.

We coupled INSeq with an existing technique [12], modified for whole root Lux-assays, to examine initial attachment to pea roots under different pH conditions. The two approaches give good agreement in 80% of cases (31 of 39) (Table 2) and this increases to >90% when additional factors are considered such as an adequate number of TA sites in the gene for confident INSeq classification. Complete agreement is not expected as the two techniques differ slightly in their experimental details. With INSeq, a mutant library is co-inoculated onto roots, while the Lux-based system assesses attachment of a homogenous population (WT or single mutant strain). These experimental differences can result in apparent discrepancies and also highlight limitations of the two techniques. RL2969 (encoding a transmembrane protein), although classified NE for attachment at pH 6.5 -7.5, showed significantly reduced attachment when used as a single inoculum in a root attachment assay (between 8% of WT at pH 6.5 and 67% at pH 7.5, Table 2). However, when co-inoculated with WT, the mutant’s inability to attach was complemented, showing that factor(s) missing in the mutant, could be provided *in trans*. A role for RL3273 in initial root attachment is also clear from the Lux-based root attachment results, despite lack of corroboration by the INSeq classification (Table 2). Despite attempts to complement mutation of RL3273 by up to a hundred-fold excess WT bacteria, we were unable to show an increase in attachment of the mutant by the presence of other bacteria. RL3273 encodes a von Willebrand factor type A (VWA) domain-containing protein, widely implicated in cell adhesion and intracellular enzyme activity in eukaryotes, and widely conserved [86]. Within rhizobia, *R. leguminosarum* has at least five [87] and *Rhizobium loti* has 12 VWA domain proteins (one magnesium chelatase and 11 uncharacterised), but *E. meliloti* has just three [86].

In agreement with previous results [12], the Lux-based assay showed that a *rapA2/C* double mutant of Rlv3841 is deficient in attachment to pea roots (Fig. 1). Genes encoding Raps (pRL100451 (*rapA2*), RL3911 (*rapB*) and RL3074 (*rapC*)) were all NE in InSeq at pH 6.5, pH 7.0 and pH 7.5. Thus, in agreement with Frederix et al [12], Rap proteins act together to facilitate attachment, and loss of one can be compensated by the presence of others.

Validation of this whole-root Lux-based assay (Table 2) represents an important development in the suite of tools available for studying early-stage root microbe interactions, without the disadvantages of using root sections, vortex and plating methods, or scintillation assays. It is the only tool to have been demonstrated as suitable for testing primary root attachment under different pH conditions.

Results from *mariner* transposon-based INSeq, Lux-based attachment assays and proteomics reveal that primary root attachment of Rlv3841 to pea roots is even more complex than previously described [3-6]. Whereas previous models have largely described a dual glucomannan/rhicadhesin system for primary root attachment under different environmental pH conditions [3-6], our work reveals a total of 115 genes whose products are specifically involved in primary attachment to plant roots, under one or more conditions in the range pH 6.5 to pH 7.5 (Fig. 3, Supplementary Table S3). However, such a list should not be considered exhaustive as we have shown the importance of RL2969, RL3273 and Raps in attachment, although this in contradiction to their INSeq classifications (Table 2).

In the final part of our work, we examined the hypothesis that rhicadhesin is a single protein required at alkaline pH for attachment using a combination of INSeq and proteomic approaches (Fig. 5, Table 4). While we confirmed that sonicated cell surface fractions do inhibit attachment at alkaline pH there were no convincing overlaps between proteins identified by proteomics and the INseq analysis. This leads us to conclude that while this fraction has attachment-inhibiting properties, it did not necessarily contain adhesins. Furthermore, INSeq analysis enabled identification of seven potential proteins; required for attachment with the necessary pH profile and of the expected gene and protein size (approx. 0.4 kb and MW respectively) (Table 4). While four of these are predicted to bind Ca^2+^ ions, none were present in the proteomic screen. While there are numerous possibilities for this, the evidence from this study is there appears to be no single specific rhicadhesin factor needed for alkaline attachment, and that using the ability to inhibit attachment is not the same as identifying a true adhesin.

In conclusion, we have shown how disruption of multiple primary attachment factors affect downstream events leading to symbiosis. It is apparent there are layers of control of gene expression, both specific and global, which would benefit from other approaches to investigate the underpinning gene networks in detail. This research has initiated a deeper understanding of how environmental pH affects rhizobial attachment to roots, a key interaction between bacteria and plants.

## Data statement

All data needed to evaluate the conclusions in this paper are present in the paper and/or Supplementary Materials.

## Acknowledgments

This work was supported by the 1379 Society, New College, University of Oxford, the Interdisciplinary Biosciences Doctoral Training Partnership, University of Oxford, the Biotechnology and Biological Sciences Research Council [grant numbers BB/N003608/1, BB/N013387/1], and the Natural Environmental Research Council [grant number NE/L501530/1]. We also wish to thank Helen Prescott and Lida Chen for their technical assistance.

## Literature Cited

1. Oldroyd GED, Downie JA. Coordinating nodule morphogenesis with Rhizobial infection in legumes. Ann Rev Plant Biol. 2008;59:519–46.

2. Poole PS, Udvardi M. Transport and Metabolism in Legume-Rhizobia Symbioses. Ann Rev Plant Biol. 2013;64:781–805.

3. Wheatley RM, Poole PS. Mechanisms of bacterial attachment to roots. FEMS Microbiol Rev. 2018;42:448–61.

4. Williams A, Wilkinson A, Krehenbrink M, Russo DM, Zorreguieta A, Downie JA. Glucomannan-mediated attachment of Rhizobium leguminosarum to pea root hairs is required for competitive nodule infection. J Bacteriol. 2008;190:4706–15.

5. Downie JA. The roles of extracellular proteins, polysaccharides and signals in the interactions of rhizobia with legume roots. FEMS Microbiol Rev. 2010;34:150–70.

6. Laus MC, Logman TJ, Lamers GE, Van Brussel AAN, Carlson RW, Kijne JW. A novel polar surface polysaccharide from Rhizobium leguminosarum binds host plant lectin. Mol Microbiol. 2006;59:1704–13.

7. Russo DM, Williams A, Edwards A, Posadas DM, Finnie C, Dankert M, et al. Proteins exported via the PrsD-PrsE type I secretion system and the acidic exopolysaccharide are involved in biofilm formation by Rhizobium leguminosarum. J Bacteriol. 2006;188:4474–86.

8. Berne C, Ducret A, Hardy GG, Brun YV. Adhesins involved in attachment to abiotic surfaces by Gram-negative bacteria. Microbiol Spec. 2015;3.

9. Ausmees N, Jacobsson K, Lindberg M. A unipolarly located, cell-surface-associated agglutinin, RapA, belongs to a family of Rhizobium-adhering proteins (Rap) in Rhizobium leguminosarum bv. trifolii. Microbiol. 2001;147:549–59.

10. Abdian PL, Caramelo JJ, Ausmees N, Zorreguieta A. RapA2 Is a calcium-binding lectin composed of two highly conserved cadherin-like domains that specifically recognize Rhizobium leguminosarum acidic exopolysaccharides. J Biol Chem. 2013;288:2893–904.

11. Mongiardini EJ, Ausmees N, Perez-Gimenez J, Althabegoiti MJ, Quelas JI, Lopez-Garcia SL, et al. The rhizobial adhesion protein RapA1 is involved in adsorption of rhizobia to plant roots but not in nodulation. FEMS Microbiol Ecol. 2008;65:279–88.

12. Frederix M, Edwards A, Swiderska A, Stanger A, Karunakaran R, Williams A, et al. Mutation of praR in Rhizobium leguminosarum enhances root biofilms, improving nodulation competitiveness by increased expression of attachment proteins. Mol Microbiol. 2014;93:464–78.

13. Ramachandran V, East AK, Karunakaran R, Downie JA, Poole PS. Adaptation of Rhizobium leguminosarum to pea, alfalfa and sugar beet rhizospheres investigated by comparative transcriptomics. Genome Biol. 2011;12:1–12.

14. Tkacz A, Poole P. Role of root microbiota in plant productivity. J Exp Bot. 2015;66:2167–75.

15. Zahran HH. Rhizobium-legume symbiosis and nitrogen fixation under severe conditions and in an arid climate. Microbiol Mol Biol Rev. 1999;63:968–89.

16. Mills KK, Bauer WD. Rhizobium attachment to clover roots. J Cell Sci. 1985;2:333–45.

17. Albareda M, Dardanelli MS, Sousa C, Megías M, Temprano F, Rodríguez-Navarro DN. Factors affecting the attachment of rhizospheric bacteria to bean and soybean roots. FEMS Microbiol Lett. 2006;259:67–73.

18. Brisset MN, Rodriguez-Palenzuela P, Burr TJ, Collmer A. Attachment, chemotaxis, and multiplication of Agrobacterium tumefaciens biovar 1 and biovar 3 on grapevine and pea. Appl Environ Microbiol. 1991;57:3178–82.

19. Johnston AWB, Beringer JE. Identification of the Rhizobium strains in pea root nodules using genetic markers. J Gen Microbiol. 1975;87:343–50.

20. Perry BJ, Yost CK. Construction of a mariner-based transposon vector for use in insertion sequence mutagenesis in the Rhizobiaceae. BMC Microbiol. 2014;14:1–11.

21. Picardeau M. Transposition of fly mariner elements into bacteria as a genetic tool for mutagenesis. Genet. 2010;138:551–8.

22. Wheatley RM, Ramachandran VK, Geddes BA, Perry BJ, Yost CK, Poole PS. Role of O2 in the Growth of Rhizobium leguminosarum bv. viciae 3841 on glucose and succinate. J Bacteriol. 2017;199:e00572–16.

23. Wheatley RM, Ford BL, Li L, Aroney STN, Knights HE, Ledermann R, et al. Lifestyle adaptations of Rhizobium from rhizosphere to symbiosis. Proc Natl Acad Sci U S A. 2020;117:23823–34.

24. Green MR, Sambrook J. Molecular Cloning Cold Spring Harbor Laboratory Press USA 2012.

25. Karunakaran R, Ramachandran VK, Seaman JC, East AK, Moushine B, Mauchline TH, et al. Transcriptomic analysis of Rhizobium leguminosarum b.v. viciae in symbiosis with host plants Pisum sativum and Vicia cracca. J Bacteriol. 2009;191:4002–14.

26. Figurski DH, Helinski DR. Replication of an origin-containing derivative of plasmid RK2 dependent on a plasmid function provided in trans. Proc Natl Acad Sci U S A. 1979;76:1648–52.

27. Martin M. Cutadapt removes adapter sequences from high-throughput sequencing reads. EMBnet. 2011;17:10–2.

28. Langmead B, Trapnell C, Pop M, Salzberg SL. Ultrafast and memory-efficient alignment of short DNA sequences to the human genome. Genome Biol. 2009;10:R25.

29. DeJesus M, Ioerger T. A Hidden Markov Model for identifying essential and growth-defect regions in bacterial genomes from transposon insertion sequencing data. BMC Bioinformat. 2013;14:303.

30. Smit G, Logman TJJ, Boerrigter METI, Kijne JW, Lugtenberg BJJ. Purification and partial characterization of the Rhizobium leguminosarum biovar viciae Ca2+-dependent adhesin, which mediates the first step in attachment of cells of the family Rhizobiaceae to plant root hair tips. J Bacteriol. 1989;171:4054–62.

31. Hu X, Dong Q, Yang J, Zhang Y. Recognizing metal and acid radical ion-binding sites by integrating ab initio modeling with template-based transferals. Bioinformat. 2016;32:3260–9.

32. Kaschani F, Gu C, Niessen S, Hoover H, Cravatt BF, van der Hoorn RA. Diversity of serine hydrolase activities of unchallenged and botrytis-infected Arabidopsis thaliana. Mol Cell Proteomics. 2009;8:1082–93.

33. Rappsilber J, Mann M, Ishihama Y. Protocol for micro-purification, enrichment, pre-fractionation and storage of peptides for proteomics using StageTips. Nature Protocols. 2007;2:1896–906.

34. Michalski A, Damoc E, Lange O, Denisov E, Nolting D, Müller M, et al. Ultra high resolution linear ion trap Orbitrap mass spectrometer (Orbitrap Elite) facilitates top down LC MS/MS and versatile peptide fragmentation modes. Mol Cell Proteomics. 2012;11:O111.013698.

35. Olsen JV, de Godoy LM, Li G, Macek B, Mortensen P, Pesch R, et al. Parts per million mass accuracy on an Orbitrap mass spectrometer via lock mass injection into a C-trap. Mol Cell Proteomics. 2005;4:2010–21.

36. Cox J, Neuhauser N, Michalski A, Scheltema RA, Olsen JV, Mann M. Andromeda: a peptide search engine integrated into the MaxQuant environment. J Proteome Res. 2011;10:1794–805.

37. Cox J, Mann M. MaxQuant enables high peptide identification rates, individualized p.p.b.-range mass accuracies and proteome-wide protein quantification. Nat Biotechnol. 2008;26:1367–72.

38. Cox J, Hein MY, Luber CA, Paron I, Nagaraj N, Mann M. Accurate proteome-wide label-free quantification by delayed normalization and maximal peptide ratio extraction, termed MaxLFQ. Mol Cell Proteomics. 2014;13:2513–26.

39. Tyanova S, Temu T, Sinitcyn P, Carlson A, Hein MY, Geiger T, et al. The Perseus computational platform for comprehensive analysis of (prote)omics data. Nat Methods. 2016;13:731–40.

40. Vizcaíno JA, Csordas A, del-Toro N, Dianes JA, Griss J, Lavidas I, et al. 2016 update of the PRIDE database and its related tools. Nucleic Acids Res. 2016;44:D447–56.

41. Schulte CCM, Ramachandran VK, Papachristodoulou A, Poole PS. Genome-Scale Metabolic Modelling of Lifestyle Changes in Rhizobium leguminosarum. 2022;7:e0097521.

42. Kearse M, Moir R, Wilson A, Stones-Havas S, Cheung M, Sturrock S, et al. Geneious Basic: an integrated and extendable desktop software platform for the organization and analysis of sequence data. Bioinformat. 2012;28:1647–9.

43. Gish W, States DJ. Identification of protein coding regions by database similarity search. Nat Genet. 1993;3:266–72.

44. Szklarczyk D, Franceschini A, Wyder S, Forslund K, Heller D, Huerta-Cepas J, et al. STRING v10: protein-protein interaction networks, integrated over the tree of life. Nucleic Acids Res. 2015;43:D447–52.

45. Yu NY, Wagner JR, Laird MR, Melli G, Rey S, Lo R, et al. PSORTb 3.0: improved protein subcellular localization prediction with refined localization subcategories and predictive capabilities for all prokaryotes. Bioinformat. 2010;26:1608–15.

46. Grant CE, Bailey TL, Noble WS. FIMO: scanning for occurrences of a given motif. Bioinformat. 2011;27:1017–8.

47. Perry BJ, Akter MS, Yost CK. The Use of Transposon Insertion Sequencing to Interrogate the Core Functional Genome of the Legume Symbiont Rhizobium leguminosarum. Front Microbiol. 2016;7:1873.

48. Becker P, Hufnagle W, Peters G, Herrmann M. Detection of differential gene expression in biofilm-forming versus planktonic populations of Staphylococcus aureus using micro-representational-difference analysis. Appl Environ Microbiol. 2001;67:2958–65.

49. Pancholi V, Fischetti VA. alpha-enolase, a novel strong plasmin(ogen) binding protein on the surface of pathogenic streptococci. J Biol Chem. 1998;273:14503–15.

50. Modun B, Williams P. The staphylococcal transferrin-binding protein is a cell wall glyceraldehyde-3-phosphate dehydrogenase. Infect Immun. 1999;67:1086–92.

51. Bao S, Chen D, Yu S, Chen H, Tan L, Hu M, et al. Characterization of triosephosphate isomerase from Mycoplasma gallisepticum. FEMS Microbiol Lett. 2015;362:fnv140.

52. Campbell GRO, Taga ME, Mistry K, Lloret J, Anderson PJ, Roth JR, et al. Sinorhizobium meliloti bluB is necessary for production of 5,6-dimethylbenzimidazole, the lower ligand of B12. Proc Natl Acad Sci U S A. 2006;103:4634–9.

53. Kumar CM, Mande SC, Mahajan G. Multiple chaperonins in bacteria--novel functions and non-canonical behaviors. Cell Stress Chaper. 2015;20:555–74.

54. Zhang Y, Cottet SE, Ealick SE. Structure of Escherichia coli AMP nucleosidase reveals similarity to nucleoside phosphorylases. Structure. 2004;12:1383–94.

55. Kumar KK, Srivastava R, Sinha VB, Michalski J, Kaper JB, Srivastava BS. recA mutations reduce adherence and colonization by classical and El Tor strains of Vibrio cholerae. Microbiol 1994;140 (Pt 5):1217-22.

56. Domínguez-Ferreras A, Pérez-Arnedo R, Becker A, Olivares J, Soto MJ, Sanjuán J. Transcriptome profiling reveals the importance of plasmid pSymB for osmoadaptation of Sinorhizobium meliloti. J Bacteriol. 2006;188:7617–25.

57. Arcus VL, McKenzie JL, Robson J, Cook GM. The PIN-domain ribonucleases and the prokaryotic VapBC toxin–antitoxin array. Prot Eng Design Select. 2010;24:33–40.

58. Puskas LG, Nagy ZB, Kelemen JZ, Ruberg S, Bodogai M, Becker A, et al. Wide-range transcriptional modulating effect of ntrR under microaerobiosis in Sinorhizobium meliloti. Mol Genet Genom. 2004;272:275–89.

59. Oke V, Rushing BG, Fisher EJ, Moghadam-Tabrizi M, Long SR. Identification of the heat-shock sigma factor RpoH and a second RpoH-like protein in Sinorhizobium meliloti. Microbiol. 2001;147:2399–408.

60. Barnett MJ, Bittner AN, Toman CJ, Oke V, Long SR. Dual RpoH sigma factors and transcriptional plasticity in a symbiotic bacterium. J Bacteriol. 2012;194:4983–94.

61. Untiet V, Karunakaran R, Kramer M, Poole P, Priefer U, Prell J. ABC Transport Is Inactivated by the PTSNtr under Potassium Limitation in Rhizobium leguminosarum 3841. PLoS One. 2013;8.

62. Tsai TI, Li ST, Liu CP, Chen KY, Shivatare SS, Lin CW, et al. An Effective Bacterial Fucosidase for Glycoprotein Remodeling. ACS Chem Biol. 2017;12:63–72.

63. Bateman A, Bycroft M. The structure of a LysM domain from E. coli membrane-bound lytic murein transglycosylase D (MltD). J Mol Biol. 2000;299:1113–9.

64. Krachler AM, Orth K. Functional characterization of the interaction between bacterial adhesin multivalent adhesion molecule 7 (MAM7) protein and its host cell ligands. J Biol Chem. 2011;286:38939–47.

65. Valentini M, Filloux A. Biofilms and Cyclic di-GMP (c-di-GMP) Signaling: Lessons from Pseudomonas aeruginosa and Other Bacteria. J Biol Chem. 2016;291:12547–55.

66. Plate L, Marletta MA. Nitric oxide modulates bacterial biofilm formation through a multicomponent cyclic-di-GMP signaling network. Mol Cell. 2012;46:449–60.

67. Schulte JE, Goulian M. The Phosphohistidine Phosphatase SixA Targets a Phosphotransferase System. Mbio. 2018;9:e01666–18.

68. Beloin C, Valle J, Latour-Lambert P, Faure P, Kzreminski M, Balestrino D, et al. Global impact of mature biofilm lifestyle on Escherichia coli K-12 gene expression. Mol Microbiol. 2004;51:659–74.

69. Minder AC, de Rudder KEE, Narberhaus F, Fischer HM, Hennecke H, Geiger O. Phosphatidylcholine levels in Bradyrhizobium japonicum membranes are critical for an efficient symbiosis with the soybean host plant. Mol Microbiol. 2001;39:1186–98.

70. Vedam V, Haynes JG, Kannenberg EL, Carlson RW, Sherrier DJ. A Rhizobium leguminosarum lipopolysaccharide lipid-A mutant induces nitrogen-fixing nodules with delayed and defective bacteroid formation. Mol Plant Microbe Interact. 2004;17:283–91.

71. Vanderlinde EM, Muszyński A, Harrison JJ, Koval SF, Foreman DL, Ceri H, et al. Rhizobium leguminosarum biovar viciae 3841, deficient in 27-hydroxyoctacosanoate-modified lipopolysaccharide, is impaired in desiccation tolerance, biofilm formation and motility. Microbiol. 2009;155:3055–69.

72. Vanderlinde EM, Harrison JJ, Muszyński A, Carlson RW, Turner RJ, Yost CK. Identification of a novel ABC transporter required for desiccation tolerance, and biofilm formation in Rhizobium leguminosarum bv. viciae 3841. FEMS Microbiol Ecol. 2010;71:327–40.

73. Yepes A, Schneider J, Mielich B, Koch G, García-Betancur JC, Ramamurthi KS, et al. The biofilm formation defect of a Bacillus subtilis flotillin-defective mutant involves the protease FtsH. Mol Microbiol. 2012;86:457–71.

74. Skagia A, Zografou C, Vezyri E, Venieraki A, Katinakis P, Dimou M. Cyclophilin PpiB is involved in motility and biofilm formation via its functional association with certain proteins. Genes Cells. 2016;21:833–51.

75. Schumann W. FtsH--a single-chain charonin? FEMS Microbiol Rev. 1999;23:1–11.

76. Kihara A, Akiyama Y, Ito K. A protease complex in the Escherichia coli plasma membrane: HflKC (HflA) forms a complex with FtsH (HflB), regulating its proteolytic activity against SecY. EMBO J. 1996;15:6122–31.

77. Mathew R, Mukherjee R, Balachandar R, Chatterji D. Deletion of the rpoZ gene, encoding the omega subunit of RNA polymerase, results in pleiotropic surface-related phenotypes in Mycobacterium smegmatis. Microbiol. 2006;152:1741–50.

78. Bhardwaj N, Syal K, Chatterji D. The role of ω-subunit of Escherichia coli RNA polymerase in stress response. Genes Cells. 2018;23:357–69.

79. Weiss A, Moore BD, Tremblay MHJ, Chaput D, Kremer A, Shaw LN. The ω Subunit Governs RNA Polymerase Stability and Transcriptional Specificity in Staphylococcus aureus. 2017;199.

80. Torres-Quesada O, Oruezabal RI, Peregrina A, Jofre E, Lloret J, Rivilla R, et al. The Sinorhizobium meliloti RNA chaperone Hfq influences central carbon metabolism and the symbiotic interaction with alfalfa. BMC Microbiol. 2010;10:-.

81. Jiang X, Zhang H, Yang J, Liu M, Feng H, Liu X, et al. Induction of gene expression in bacteria at optimal growth temperatures. Appl Microbiol Biotechnol. 2013;97:5423–31.

82. Griffin JE, Gawronski JD, DeJesus MA, Ioerger TR, Akerley BJ, Sassetti CM. High-Resolution Phenotypic Profiling Defines Genes Essential for Mycobacterial Growth and Cholesterol Catabolism. PLoS Path. 2011;7:e1002251.

83. Zhang YJ, Ioerger TR, Huttenhower C, Long JE, Sassetti CM, Sacchettini JC, et al. Global assessment of genomic regions required for growth in Mycobacterium tuberculosis. Plos Pathog. 2012;8:e1002946.

84. Fox MA, Karunakaran R, Leonard ME, Mouhsine B, Williams A, East AK, et al. Characterization of the quaternary amine transporters of Rhizobium leguminosarum bv. viciae 3841. FEMS Microbiol Lett. 2008;287:212–20.

85. diCenzo GC, Muhammed Z, Østerås M, O’Brien SAP, Finan TM. A Key Regulator of the Glycolytic and Gluconeogenic Central Metabolic Pathways in Sinorhizobium meliloti. Genet. 2017;207:961–74.

86. Whittaker CA, Hynes RO. Distribution and evolution of von Willebrand/integrin A domains: widely dispersed domains with roles in cell adhesion and elsewhere. Mol Biol Cell. 2002;13:3369–87.

87. Young JP, Crossman LC, Johnston AW, Thomson NR, Ghazoui ZF, Hull KH, et al. The genome of Rhizobium leguminosarum has recognizable core and accessory components. Genome Biol. 2006;7:R34.

